# c-Rel drives pancreatic cancer metastasis through Fibronectin-Integrin signaling-induced isolation stress resistance and EMT activation

**DOI:** 10.1101/2025.01.29.635445

**Authors:** D. Bakırdöğen, K. Görgülü, J. Xin, S. Alcalá, L. Ruiz-Cañas, K. Frank, N. Wu, K. N. Diakopoulos, C. Dai, H. Öztürk, D. Demircioğlu, K. Peschke, R. Ranjan, F. Fusco, J. Martinez-Useros, M. J. Fernandez-Aceñero, N. F. Chhabra, J.C. López-Gil, J. Ai, D. A. Ruess, E. Kaya-Aksoy, K. Steiger, F. Schmidt, L. Kohlmann, A. Berninger, R. M. Schmid, M. Reichert, M. Adli, M. Lesina, B. Sainz, H. Algül

**Affiliations:** Comprehensive Cancer Center Munich CCCM, Technische Universität München, Munich, Germany; Cancer Stem Cells and Fibroinflammatory Microenvironment Group, Cancer Department, Instituto de Investigaciones Biomédicas (IIBM) Sols-Morreale CSIC-UAM, Madrid, Spain; Biomarkers and Personalized Approach to Cancer Group (BIOPAC), Area 3 Cancer, Instituto Ramón y Cajal de Investigación Sanitaria (IRYCIS), 28049, Madrid, Spain; Centro de Investigación Biomédica en Red, Área Cáncer, CIBERONC, ISCIII, Madrid, Spain; Robert Lurie Comprehensive Cancer Center, Department of Obstetrics and Gynecology, Feinberg School of Medicine at Northwestern University, Chicago, IL, USA; Department of Oncological Sciences, Icahn School of Medicine at Mount Sinai, New York, NY, 10029; Tisch Cancer Institute, Icahn School of Medicine at Mount Sinai, New York, NY, 10029; Bioinformatics for Next Generation Sequencing (BiNGS) core, Icahn School of Medicine at Mount Sinai, New York, NY, 10029, USA; Translational Pancreatic Cancer Research Center, TUM School of Medicine and Health, Department of Clinical Medicine – Clinical Department for Internal Medicine II, University Medical Center, Technical University of Munich; TUM School of Medicine and Health, Department of Clinical Medicine – Clinical Department for Internal Medicine II, University Medical Center, Technical University of Munich; Center for Protein Assemblies (CPA), Technical University of Munich, Germany; Center for Organoid Systems (COS), Technische Universität München, Germany; German Cancer Consortium (DKTK), partner site Munich, Munich, Germany; Bavarian Cancer Research Center (BZKF), Munich, Germany; Institute of Pathology, School of Medicine, Technical University of Munich, Munich, Germany; Translational Oncology Division, Oncohealth Institute, Fundacion Jiménez Díaz University Hospital, 28040 Madrid, Spain; Area of Physiology, Department of Basic Health Sciences, Faculty of Health Sciences, Rey Juan Carlos University, 28922 Madrid, Spain; Pathology Department, Clinico San Carlos University Hospital, 28040 Madrid, Spain; Department of Gastroenterology, The First Affiliated Hospital of Nanchang University, Nanchang, Jiangxi province, China; Department of General and Visceral Surgery, Center for Surgery, Medical Center University of Freiburg, Freiburg, Germany; German Cancer Consortium (DKTK), Partner Site Freiburg and German Cancer Research Center (DKFZ), Heidelberg, Germany

**Author notes:** CORRESPONDANCE Address correspondence to: Hana Algül, MD, MPH, and Derya Bakirdögen, Dr. rer. Nat. Comprehensive Cancer Center, Klinikum der Technischen Universität München, Technische Universität München, Ismaninger Str. 22, 81675 Munich, Germany.

## Abstract

Pancreatic ductal adenocarcinoma remains one of the deadliest malignancies, with limited treatment options and a high recurrence rate. Recurrence happens often with metastasis, for which cancer cells must adapt to isolation stress to successfully colonize distant organs. While the fibronectin-integrin axis has been implicated in this adaptation, its regulatory mechanisms require further elaboration. Here, we identify c-Rel as an oncogenic driver in PDAC, promoting epithelial-to-mesenchymal transition (EMT) plasticity, extracellular matrix (ECM) remodeling, and resistance to isolation stress. Mechanistically, c-Rel directly regulates fibronectin (*Fn1*) and CD61 (*itgb3*) transcription, enhancing cellular plasticity and survival under anchorage-independent conditions. Fibronectin is not essential for EMT, but its absence significantly impairs metastatic colonization, highlighting a tumor-autonomous role for FN1 in isolation stress adaptation. These findings establish c-Rel as a key regulator of PDAC metastasis by controlling circulating tumor cell (CTC) niche and survival, suggesting that targeting the c-Rel–fibronectin–integrin axis could provide new therapeutic strategies to mitigate disease progression and recurrence.

## INTRODUCTION

Pancreatic cancer (PDAC) is the third leading cause of cancer-related deaths, with a 13% 5-year survival rate in the United States ^1^. Although the overall cancer mortality for all cancers trend to decline, pancreatic cancer incidence increases ^1^. Projections for both 2030 and 2040 suggest that pancreatic cancer-related death will be the second in place, trailing behind lung cancer ^2,3^. Nevertheless, treatment options are still limited ^4^. Patients with resected PDAC tumors recur rather fast (∼75%), owing to circulating tumor cells (CTCs) and already established distant micro-metastases (∼60%) ^5,6^. Although adjuvant chemotherapy may reduce metastatic recurrence, more effective targeted approaches are required.

Within the metastatic cascade, cancer cells once in circulation have the adaptability to overcome isolation stress ^7^. Tolerance to isolation stress induced by limiting survival conditions including nutrient stress, hypoxia, oxidative stress, or detachment from the extracellular matrix (ECM), can enhance cancer cell resilience. These cancer cells can demonstrate boosted stemness, tumor-initiating capacity, and metastasis. Recently, an enriched fibronectin-integrin axis has been shown to increase isolation stress tolerance in PDAC ^8^. In parallel, multiple studies identified various subclusters of pancreatic CTCs with the expression of ECM-associated genes ^9–12^. An expanded understanding of the signaling networks regulating ECM-induced isolation stress tolerance is required to develop novel therapies against metastatic PDAC recurrence.

With its multiple components, NF-κB is a ubiquitous signaling pathway active in various physiological and pathophysiological processes ^13^. Due to the technical limitations, NF-κB signaling is long thought to show binary activation (on/off). However, recent advances in techniques, enabling single-cell resolution for analysis, have allowed multiple quantitative features of NF-κB signaling dynamics to be revealed ^14^. Despite all these technical advances, the functional roles of the individual NF-κB components in pancreatic pathophysiology remain sparse.

Of the five NF-κB transcription factors, RelA, RelB, and c-Rel contain a transactivation domain (TAD), whose heterodimers can induce transcriptional activation of target genes. Previously, our group demonstrated a context-specific function of the canonical NF-κB transcription factor RelA (p65) in PDAC ^15^. Interestingly, during carcinogenesis, RelA assumes a tumor suppressor role due to its involvement in senescence activation in cancer precursors. However, once the senescence checkpoint is exceeded, the oncogenic function of RelA becomes rather operative. Additionally, the absence of the noncanonical NF-κB transcription factor RelB has been shown to decelerate pancreatic carcinogenesis ^16^.

In contrast, REL proto-oncogene, NF-kB subunit (c-Rel) is not studied meticulously in solid tumors, including PDAC. Previous studies have reported both tumor suppressor ^17–19^ and oncogenic ^20–23^ roles for c-Rel in cancer. Additional studies also highlight c-Rel’s involvement in TNF-related apoptosis-inducing ligand (TRAIL) resistance ^24^, epithelial to mesenchymal transition (EMT) ^25–28^, cancer stem cell characteristics (CSCness) ^28,29^, and immunomodulatory effects ^30–33^ in various cancers.

Herein, utilizing genetically engineered mouse models (GEMMs) of PDAC, we uncover a predominantly oncogenic role for heightened c-Rel protein activity. We demonstrate that c-Rel acts as a negative prognostic factor in PDAC survival, where tumors exhibiting elevated c-Rel expression are characterized by an undifferentiated morphology with EMT plasticity and contractility, and ECM remodeling. Specifically, tumors with high c-Rel levels have an enhanced Fibronectin (FN1) – Itgb3 signaling, allowing them to have higher tolerance in isolation stress and metastatic homing. Overall, our findings suggest a novel oncogenic role for c-Rel in PDAC.

## RESULTS

### 1) c-Rel is expressed in both human and murine PDAC

To assess c-Rel’s ubiquity in PDAC, we analyzed various human and mouse samples. c-Rel is expressed ubiquitously in a panel of cell lines isolated from human and mouse PDAC (SF1A). Of note, mouse PDAC cell lines are isolated from GEMMs (CK: *Ptf1a-Cre Kras^G12D^*, CKP: *Trp53^f/f^ Ptf1a-Cre Kras^G12D^*). IHC analysis revealed c-Rel expression in cancer cells, other TME components, and pre-malignant PanIN lesions in mouse tumors, with minimal signal in the acini (SF1B).

Investigating the prognostic impact of *REL* mRNA in PDAC, we analyzed the TCGA, CPTAC-3, and Bailey datasets via the pdacR online tool ^34^. We categorized patients into high vs. low *REL* expression groups, using cutoff values of 0.5, 0.65, and 0.75 to compare top-median, top-tertile, and top-quartile respectively. Although almost none of the findings were statistically significant, a trend for poorer prognosis in high *REL* expression patients was observed in the TCGA and Bailey cohorts (SF1C). In a cohort of resected human PDAC patients, c-Rel protein expression was noted in 149 out of 174 patients (F1A). The histogram plot indicated that most tumor samples express low levels of c-Rel, while a patient subset had very high expression (F1A). Moreover, approximately half of the patients showed c-Rel’s nuclear localization (F1A). However, neither H-score nor nuclear localization-based stratification indicated prognostic impact in the disease-specific overall survival and progression-free survival in the tumor patients (SF1D).

Previously, we reported a reduction in *NFKB_SIGNALING* signature in human PDAC spheroids under non-adherent conditions compared to their adherent counterparts ^35^. On the contrary, *REL* mRNA level is increased, unlike the NF-κB signature in the analyzed patient cell lines (F1B). In parallel, among the NF-κB proteins, only c-Rel and RelA showed a consistent increase in protein expression under spheroid culture (F1C). These results highlight the importance of c-Rel under non-adherent conditions.

Under the basal adherent culture conditions, c-Rel was mostly present in the cytoplasm, while canonical NF-κB inducer TNF-α quickly facilitated its nuclear translocation (SF1E). Of note, only a 20-minute treatment with nuclear export inhibitor Leptomycin-B was enough to accumulate c-Rel in the nucleus (SF1E). Co-immunoprecipitation analysis revealed c-Rel heterodimerization with RelA (SF1F). Moreover, although the total c-Rel levels showed a very mild reduction in total expression (SF1G), its nuclear translocation is increased in RelA knock-out CK (CKA) and CKP cells (CKPA) (SF1H), suggesting compensatory mechanisms in between. Overall, these findings indicate the involvement of c-Rel in the canonical NF-κB signaling in cooperation with RelA in PDAC.

### 2) Higher c-Rel levels drive partial-EMT and activated stroma in PDAC

To study c-Rel function more closely in PDAC using GEMMs, we generated compound mutant mice by crossing CKP mice with either a *Rel*^f/f^ knock-out model (CCKP) or a *GFP-Rel^f/+^* overexpression model (GCKP) (F1D). Of note, all the mice are screened for possible recombination of the *floxed* alleles in the body, since *Ptf1a-Cre* can cause recombination in the paternal germline with high efficiency ^36^. In these models, c-Rel expression inversely correlated with survival: GCKP had a shorter lifespan with more proliferation (F1F), while CCKP survived longer with less proliferation (F1E). In particular, GCKP tumors were larger (F1G) and exhibited higher tumor grade (F1H) and undifferentiated solid growth pattern (F1I), as blindly assessed by a pathologist.

Based on the tumor histopathology, we analyzed the EMT and its plasticity. IHC stainings (F2A), immunoblots (F2B), and RT-qPCRs (SF2A) of bulk tumor tissues revealed a decrease in epithelial markers (E-cadherin, CK-19, β-catenin) along with an increase in mesenchymal markers (Vimentin, Zeb1, Zeb2, Snai2) with a modest reduction in *Snai1* at the bulk transcriptional level. Cancer cells isolated from these mouse tumors showed greater morphological scattering *in vitro* (F2C) and in organoid cultures in matrigel (F2D). Immunoblot analyses indicated decreased E-cadherin and increased Zeb1, Vimentin, and N-Cadherin (an additional band is observed only in GCKP) in the isolated cancer cells (F2E). GCKP cells reduced surface E-cadherin expression (F2F) and displayed intracellular E-cadherin speckles aligning with partial-EMT characteristics (F2G) ^37^. The EMT difference observed between CCKP and CKP tumors *in vivo* was not evident *in vitro* under basal conditions, but the TGFβ treatment potentiated it. Under TGFβ treatment, CCKP cells resisted surface E-cadherin reduction more than CKP cells (SF2B). Multiple previously published EMT-related signatures revealed significant positive correlations with *REL* mRNA expression in PDAC samples as analyzed in the EMTome database (SF2C) ^38^. Moreover, additional PDAC patient datasets are divided from the median based on *REL* expression to be analyzed for *HALLMARK_EMT* signature, which displayed significant enrichment in *REL* high samples (SF2D). In parallel, increased c-Rel expression enhanced cancer cell contractility (F2H), with more Src and Stat3 phosphorylation, but without a difference in p-FAK (F2I). Nevertheless, the GCKP cells were more sensitive to dasatinib (Src-i), but not to defactinib (FAK-i) (SF2E).

Further histopathological examination disclosed notable changes in the ECM. Sirius-red staining quantitatively showed less collagen deposition in GCKP tumors (F3A). Conversely, FN1 expression, particularly elevated in GCKP tumors, was confirmed through IHC (F3B) and immunoblot analyses (F3C). In CCKP and CKP tumors, FN1 primarily surrounded collective tubular-cribriform structures, whereas, in GCKP tumors, it individually encircled undifferentiated cancer cells, as evidenced in the representative images of GCKP-1 (F3B). Intriguingly, some cancer cells displayed a perinuclear FN1 pattern as exemplified in GCKP-2 (F3B). ELISA suggested a significant correlation between serum FN1 and c-Rel levels in tumor-bearing mice (F3D). Additionally, increased c-Rel expression enhanced FN1 responsiveness, as GCKP cells attached to FN1 quicker (F3E).

To corroborate our findings, we performed RNA-seq analysis on the isolated CCKP, CKP, and GCKP cancer cells, with pairwise comparisons for GCKP vs CKP, GCKP vs CCKP, and CCKP vs CKP samples (F3F). Heterogeneity in cancer is the norm, as exemplified in the PCA analysis of the individual cancer cells (F3G). Still, cells with the same genotypic background grouped (F3G). To our surprise, there were no common genes that are significantly upregulated on GCKP and downregulated in CCKP (except *Rel*), or vice versa (F3H). On the other hand, 12 and 9 genes were up- and downregulated respectively in GCKP and CCKP cells compared to CKP (F3H).

We attributed the absence of common genes in the c-Rel overexpression vs knock-out setting to the differential regulation of NF-κB proteins. Therefore, we checked the total (SF3A) and nuclear expression (SF3B) for some of the targets. Among the targets, we saw a notable and gradual increase in IKBα and IKBβ proteins from knock-out CCKP to overexpression GCKP cells. There was no expression change in RelA and RelB proteins. Although p105 expression is increased, p50 levels didn’t change in GCKP samples. Under the basal condition, nuclear translocation of RelA is decreased only in CCKP cells, while RelB was reduced in both CCKP and GCKP cells. The lack of consistency in total expression and nuclear localization of most proteins highlights additional cross-talk mechanisms between them. Overall, the lack of commonly regulated genes observed in the c-Rel knock-out and overexpression setting can be attributed to the multi-level organization of variable NF-κB activation.

In line with our results, GSEA analysis revealed multiple enrichments related to EMT and ECM remodeling (F3I). Additionally, *Rel* expression correlated significantly with Moffitt’s activated stroma signatures in multiple human PDAC transcriptomics datasets (SF3C). Recently, a detailed molecular analysis for PDAC patients revealed that tumors could possess multiple stromal subtypes (subTME) ^39^. Tumors with reactive subTMEs have higher c-Rel and FN1 protein expression in tumor cells and stroma, respectively (SF3D). In parallel to these findings, tumor cell expression of c-Rel protein showed a positive correlation with FN1 presence in the stroma (SF3E). Notably, integrin signaling associated with FN1 binding was also enriched in PID (Pathway interaction database) in GCKP cells (SF3F). Further validation through GSEA analysis on human PDAC bulk transcriptomics datasets resulted in significant ECM-Receptor interaction enrichment, particularly involving integrins (SF3G). Finally, we analyzed 2 human PDAC scRNAseq datasets, each pooled with multiple patient samples. As expected, *REL* expression is mostly observed in immune cells like myeloid cells, T/NK cells, and mast cells (SF3H). The cell types are distinguished based on the expression of target genes given in the tabular results (SF3H). The analysis underscored that epithelial cells with a high REL fraction had a bigger fraction of cells expressing multiple integrin family transcripts along with more expression (SF3H). Collectively, these results indicate that c-Rel regulates EMT and ECM remodeling, possibly involving FN1-integrin signaling in PDAC.

### 3) c-Rel enhances isolation stress adaptation of PDAC cells

c-Rel expression is increased under non-adherent conditions in patient PDAC cells (F1B-C). Based on these findings, we conducted multiple assays with the cells to measure their ability to resist isolation stress. GCKP cells demonstrated more colony-forming frequency, as quantified by their ability to form organoids in Matrigel (F4A) and formed larger spheroids under adherence-free conditions (F4B). Based on these results, we tested the surface expression of predefined stemness-associated markers, including EpCAM, Cxcr4, CD44, CD133, Sca1, and CD61 (integrin β3) in adherent (SF4A) and non-adherent serum-free spheroid cultures (F4C). To our surprise, all the markers, either individually or double-positive (with EpCAM+) showed a reduction in GCKP cells, except for CD61 in spheroid cultures (F4C). In parallel, CD61 expression is increased with c-Rel levels in primary tumors (F4D). Overall, these results suggest that c-Rel is involved in CD61-dependent cell survival under isolation stress.

### 4) Ectopically expressed c-Rel increases EMT, ECM remodeling, and resistance to isolation stress of cancer cells

To validate our findings regarding c-Rel’s regulatory role in multiple aspects of PDAC behavior, we utilized the Auxin-inducible degron 2 (AID2) system (SF4B) ^40^. AID2 system allows users to rapidly knock down their protein of interest after an auxin analog (5phIAA) treatment. In two CCKP cell lines (A2929 and A2770), we overexpressed c-Rel with both GFP and AID tags (SF4C), enabling rapid degradation of c-Rel within 2 hours up on 5phIAA treatment (SF4D). Compared to the mock-transfected cells, c-Rel-pAY15 overexpression induced a scattered cellular morphology with reduced surface E-cadherin expression (F4E). Spheroid size also increased in c-Rel-pAY15 cells, and 5phIAA-induced c-Rel degradation reversed and rescued the phenotype (F4F). Moreover, the surface expression of CD61 followed a similar trend (F4G). Notably, EpCAM and CD133 surface expression increased after c-Rel degradation (SF4E).

To assess c-Rel’s direct transcriptional regulation in the associated phenotypes, we performed a CUT&RUN experiment with the degron cells ^41,42^. CUT&RUN is an alternative to Ch-IP, where antibody binding to target protein allows micrococcal nuclease recruitment and DNA cleavage, releasing target-protein DNA complexes. c-Rel’s nuclear accumulation is achieved with the nuclear export inhibitor leptomycin-B (F5A). The peaks indicated broad c-Rel binding on intergenic, intragenic, and promoter regions (F5B). Analysis of *FN1* and *Itgb3* genes suggested c-Rel binding to their promoter regions (F5B). mRNA levels of *FN1* and several RGD-integrins (*Itgb3, ItgaV, Itga5*), assessed by RT-qPCR, escalated in c-Rel-overexpression cells, especially under 3D spheroid culture conditions, and reverted to lower levels following 5phIAA induced c-Rel degradation (F5C). Therefore, we performed an RNA-seq analysis on degron cells under spheroid cultures. In parallel to the previous findings, cells overexpressing c-Rel had significant upregulation in *FN1* and *Itgb3* mRNA levels (F5B). Moreover, the over-representation analysis performed with Enrichr ^43–45^ revealed enrichments for EMT, Itgb3 signaling, and ECM modeling pathways, which were rescued by degron stimulation (F5D). The EMT regulators Snai1, Snai2, Tgfb1, and Zeb2 were also among the c-Rel DNA-binding targets (SF5A). Of note, some genes where c-Rel occupied the promoters were downregulated in the transcriptomics analyses (SF5B). Further overrepresentation analysis indicated significant downregulations for various metabolic pathways when c-Rel is overexpressed (SF5C). Overall, these results indicate c-Rel’s direct regulation in EMT and FN-integrin signaling.

### 5) Fibronectin depletion in cancer cells doesn’t alter EMT and survival

Building on the results, indicating c-Rel’s positive regulation of the FN1-integrin axis both in vivo and in vitro, we hypothesized that FN1 absence in cancer cells might counteract the GCKP-associated phenotypes. We generated compound mutant mice by crossing *the Fn1^f/f^* model with CKP and GCKP mice (F6A) to test this. Although the reduced overall survival (OS) in GCKP mice was not restored to longer CKP durations (F6B), tumor weight reverted to CKP levels (F6C). There was also no difference between the survivals of CKP and FNCKP mice (F6B). Of note, we validated the FN1 reduction in tumor tissues by RT-qPCR (SF6A), immunoblots (SF6B), and IHC (SF6C). All the FN1 isoforms tested showed a decrease in the FN1 knock-out settings (SF6A). Although FN1 was depleted in cancer cells, it remained abundantly in ECM (SF6C). Moreover, FNGCKP tumors still maintained high-grade tumors with major solid growth patterns similar to GCKP tumors (F6D). FNCKP tumors had higher tumor grade compared to CKP, although their growth pattern was not different (F6D). *FN1* depletion in cancer cells didn’t rescue EMT in vivo, neither in CKP nor GCKP tumors. Epithelial identity associated CK19, E-Cadherin, and β-Catenin and mesenchymal protein vimentin expressions didn’t change in FNCKP and FNGCKP samples compared to their controls CKP and GCKP respectively (F6E). To eliminate the FN1 assistance from stromal sources during EMT identity *in vivo*, we analyzed the cancer cells *in vitro*. The isolated cancer cells are validated for their lack of FN1 and overexpression of c-Rel by immunoblot (F6F). The absence of FN1 rescued the scattered morphology observed in neither CKP nor GCKP cells (F6G). FNCKP cells show a consistent protein expression change compared to CKP in neither of the markers checked (F6H). On the other hand, FNGCKP cells slightly increased their E-cadherin with a reduction in vimentin expression compared to GCKP cells (F6F). However, the surface expression of E-cadherin didn’t change in the CKP-FNCKP and GCKP-FNGCKP comparison (F6H). These results indicate that FN1 is not essential for EMT, and its absence can’t rescue the survival and EMT phenotypes observed in GCKP tumors.

### 6) FN1 is necessary for c-Rel-induced isolation stress resistance of cancer cells

We wanted to test whether FN1 is required to maintain CD61-associated anchorage-independent growth in GCKP cells. The colony forming frequency in matrigel displayed no change in FNCKP cells compared to CKP control (F7A). On the other hand, higher colony-forming frequency is rescued in GCKP cells after FN1 deletion (F7A). Higher average spheroid size under adherent-free conditions is also reduced in FNGCKP cells compared to GCKP (F7B).

Intriguingly, neither plasma FN1 nor cellular FN1-EDB in various concentrations rescued the spheroid size in FNGCKP cells (F7C). In line with the previous findings, in FNGCKP spheroids, the surface expression of CD61 is reduced relative to GCKP, and supplementing plasma FN1 again couldn’t rescue the phenotype (F7D). Moreover, although not significant, expressions of the other conventional markers are increased (F7D). Our results indicate the involvement of c-Rel in cellular fitness during isolation stress, an important concept in the metastatic cascade. To assess the importance of the c-Rel mediated integrin-FN1 signaling axis in the metastatic cascade, we utilized IV injection of the cancer cells followed by lung histopathology analysis. As expected, both relative metastatic colony area in the lungs and relative lung weights are increased in GCKP cells (F7E). Moreover, the absence of FN1 in GCKP cells rescued the metastatic phenotype to CKP levels. These results indicate that the c-Rel-FN1-integrin signaling axis is an important regulator of the metastatic cascade.

## DISCUSSION

Herein, we propose a mechanism where c-Rel is involved in multiple oncogenic facets, collectively regulating the survival and metastatic behavior of PDAC (F8). By directly regulating the expression of EMT-related transcription factors, c-Rel inflects EMT plasticity in PDAC. With the transcriptional regulation of key ECM structural and modifier proteins, c-Rel finetunes ECM remodeling. Finally, by directly regulating FN1 and integrin-β3 expression, c-Rel modulates anchorage-independent growth and metastatic homing of PDAC.

Although the experimental evidence suggests the importance of c-Rel function in PDAC prognosis, survival analyses obtained from TMAs or human PDAC transcriptomics don’t reveal such a profound consequence. The kinetic and dynamic heterogeneity in c-Rel nuclear translocation and transcriptional activation can shadow the probable survival difference in patient cohorts ^14^. Indeed, only a short-term inhibition of nuclear export via leptomycin-B was enough to accumulate c-Rel majorly in the nucleus, indicating a very dynamic nuclear translocation (SF1E, SF5A). Therefore, a snapshot of neither c-Rel levels nor its nuclear localization obtained from a single timepoint point can be suitable to assess c-Rel’s potential contribution to patient survival.

The phenotypic difference observed in c-Rel overexpressing tumors were also not always negatively overlapped in c-Rel knock-out samples. Although c-Rel’s nuclear localization was higher in RelA knock-outs, RelA’s localization was less in c-Rel knock-out cells. However, this doesn’t necessarily mean that RelA is less functional as a transcription factor in c-Rel knock-out cells. Variables like ligand type, temporal control of stimulation, nuclear translocation oscillation dynamics, the downstream gene targets and transcriptional co-regulatory proteins must be further elaborated to have an understanding for the crosstalk between c-Rel and RelA. Nevertheless, evidence suggests both redundant and nonredundant functions of c-Rel, in some, finetuning RelA transcriptional activation in a ligand-specific manner during inflammation ^46–51^. Such an interplay may also be evident in epithelial cell behavior in PDAC, hindering the phenotypic difference observed in the c-Rel knockout setting.

Cellular plasticity and niche interaction are among the emerging hallmarks of cancer cell stemness ^52^. Heterogeneity in cellular plasticity and niche interaction may impact CSC subtypes. For example, Lgr5+ colorectal CSCs have the ability to self-renew and differentiate into Lgr5-invasive and metastatic cancer cells ^53^. Interestingly, no Lgr5+ CSCs were detected as CTC in circulation. Moreover, the Lgr5-CTCs had no change in the conventional CSC marker expression like CD133 or CD44. Lack of CSC niche may force them to survive through alternative mechanisms. Once colonized into liver, owing to their plasticity, Lgr5-cells can partially differentiate back into Lgr5+ CSCs. A similarity has also been reported in skin cancer. Epcam-tumor cells have higher tumor-initiating and metastatic capacity compared to the Epcam+ ones ^54^. Interestingly, Epcam+ cells were enriched for epithelial signature and CSC markers like CD133, Sox2 or Aldh1. The Epcam-counterpart was enriched for EMT and ECM including FN1. Epcam-mesenchymal-like cells in skin and mammary cancer were shown to possess a grade of EMT states with the combination of CD106, CD51, and CD61 surface expression ^55^. Most of these cells in the intermediate states have comparable TIC capacity, while their invasiveness and metastasis differed. Although we majorly tested the surface expression of CD61, transcriptomics data indicates a c-Rel-dependent increase in all of these markers. In parallel, conventional CSC surface expression markers mostly showed a reduced expression pattern or no change in GCKP. Therefore, we suggest a model where c-Rel induces a tumor quasi-mesenchymal CSC subtype ^56^ with EMT plasticity in PDAC, increasing its both tumor initiating and metastatic capacity.

Previously, CD61 was shown to be an upstream inducer of c-Rel activation in erlotinib-resistant CSCs ^29^. Adding on these findings, FN1-CD61 expression under the control of c-Rel activation highlights a positive feedback loop. Moreover, CD61+ CTCs were identified as a sub-cluster in PDAC CTCs ^10^. Additionally, during isolation stress CD61/CD51 and CD29/CD49e are associated with resistance to cell death ^8,57^. Among these targets, not only CD61 and CD51 but also CD49e showed increased expression during anchorage-independent growth in transcriptomics. Therefore, we suggest a central role for c-Rel transcriptional activation in the resilience of CTCs before metastatic colonization under anchorage-independent growth.

The absence of FN1 in the GCKP tumors can’t rescue survival, which might be due to the abundance of FN1 in ECM coming from stromal cells in the primary tumors. However, cells isolated from these tumors display impaired metastatic colonization. These results hint at the tumor autonomous redundancy for FN1 production in primary tumors. However, in circulating tumor cells with metastatic potential, FN1-coupled CD61 surface expression is important. Moreover, intracellular production of FN1 stimulates anchorage-independent growth under isolation stress. Reduction of CD61 surface expression is also coupled with FN1 absence. The absence of CD61 expression on the cell surface could be due to two reasons. Differences in the post-translational modifications (e.g. citrullination ^58^) in supplemented soluble FN1 in spheroid medium might result in an inability to rescue the reduced spheroid formation in FN1 knock-out cells. Secondly, multiple integrins were shown to be shuttled to the cell surface in a fibronectin-coupled manner ^59,60^. This shuttling may also regulate FN1 fibrillogenesis and assembly at the ECM ^61,62^. Therefore, we suggest that integrin trafficking, related to the anchorage-independent growth is affected by FN1.

Recently, c-Rel has been identified by many studies as a checkpoint regulator of tumor-promoting myeloid and Treg function, along with fibrosis ^30–32^. Our model proposes a therapeutic rationale for targeting c-Rel in solid tumors as a stratification marker of isolation stress during metastasis and targeting multiple aspects of tumor biology.

**Figure 1:**
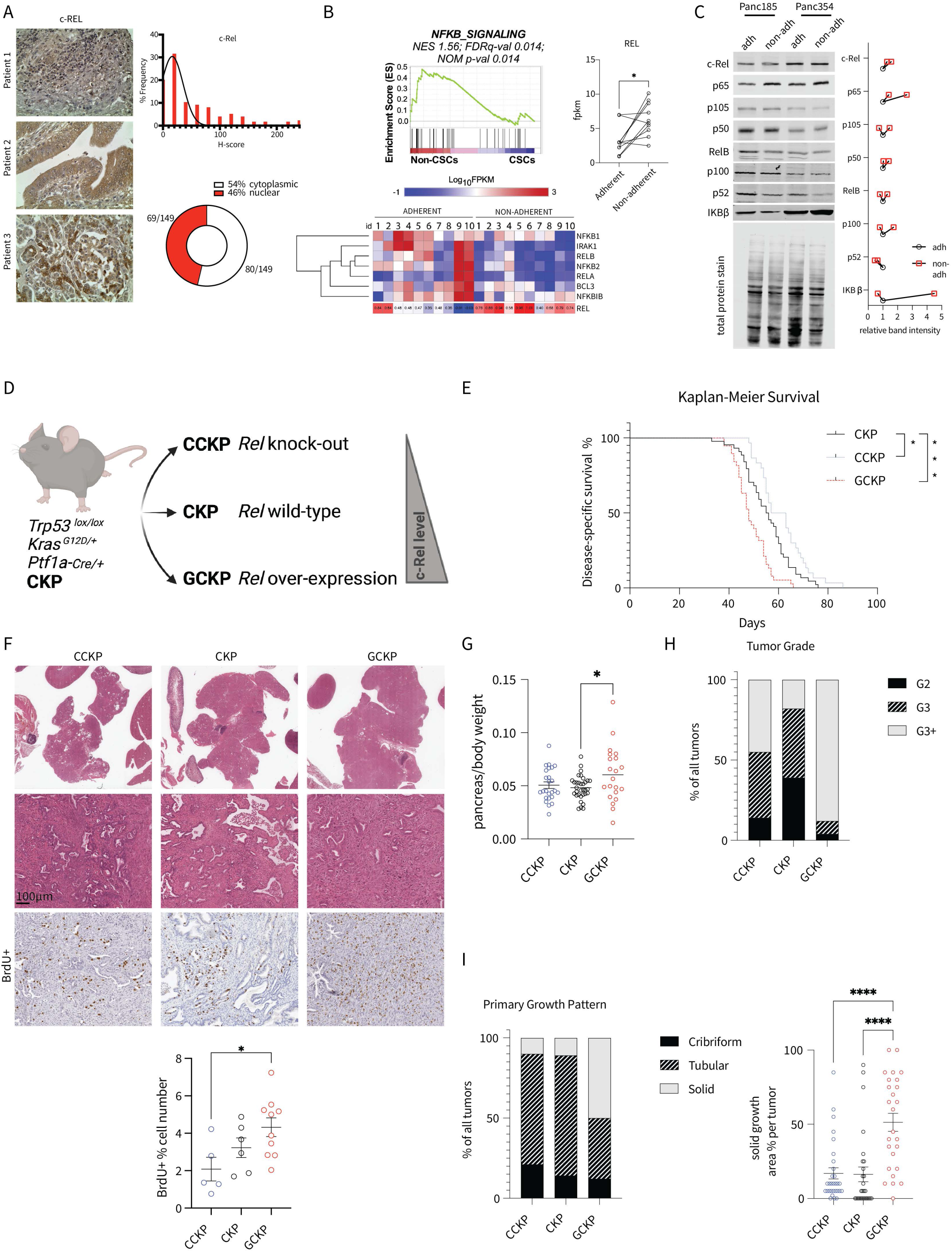
A) Representative c-Rel IHC images from patient PDAC samples. The h-score distribution is shown in the histogram plot along with the chart displaying the nuclear localization percentage. B) GSEA for human PDAC cell lines cultured under adherent vs non-adherent conditions. *REL* mRNA fpkm values are compared with paired t-test. C) Western blot analyses for patient cell lines, cultured under adherent and non-adherent conditions. The quantification is normalized to the total protein stain signal. D) Illustration demonstrating the compound mutant mice generation with *Rel* knock-out (CCKP), wild-type (WT) (CKP), and overexpression (GCKP). E) Kaplan-Meier survival curves for CCKP (n=30), CKP (n=44), and GCKP (n=38) mice. Statistical analysis was performed with the Log-rank (Mantel-Cox) test. F) Representative histology sections for H&E and BrdU IHC stainings. Statistics for the percentage of BrdU+ cells (p=*). G) Pancreas weight/body weight comparison (p=*) H, I) Chi-square test (p=****) is used for tumor grade and primary growth pattern analysis. For the solid growth area % per tumor ordinary one-way ANOVA (p=****) and Tukey’s multiple comparisons test were used. The number of tumor mice analyzed is as given: CCKP n= 29, CKP n=28, GCKP n=26. Unless otherwise indicated, for all analyses ordinary one-way ANOVA test is used. ANOVA p values are given in the figure legends. Tukey’s multiple comparison tests are shown in the figures.

**Figure 2:**
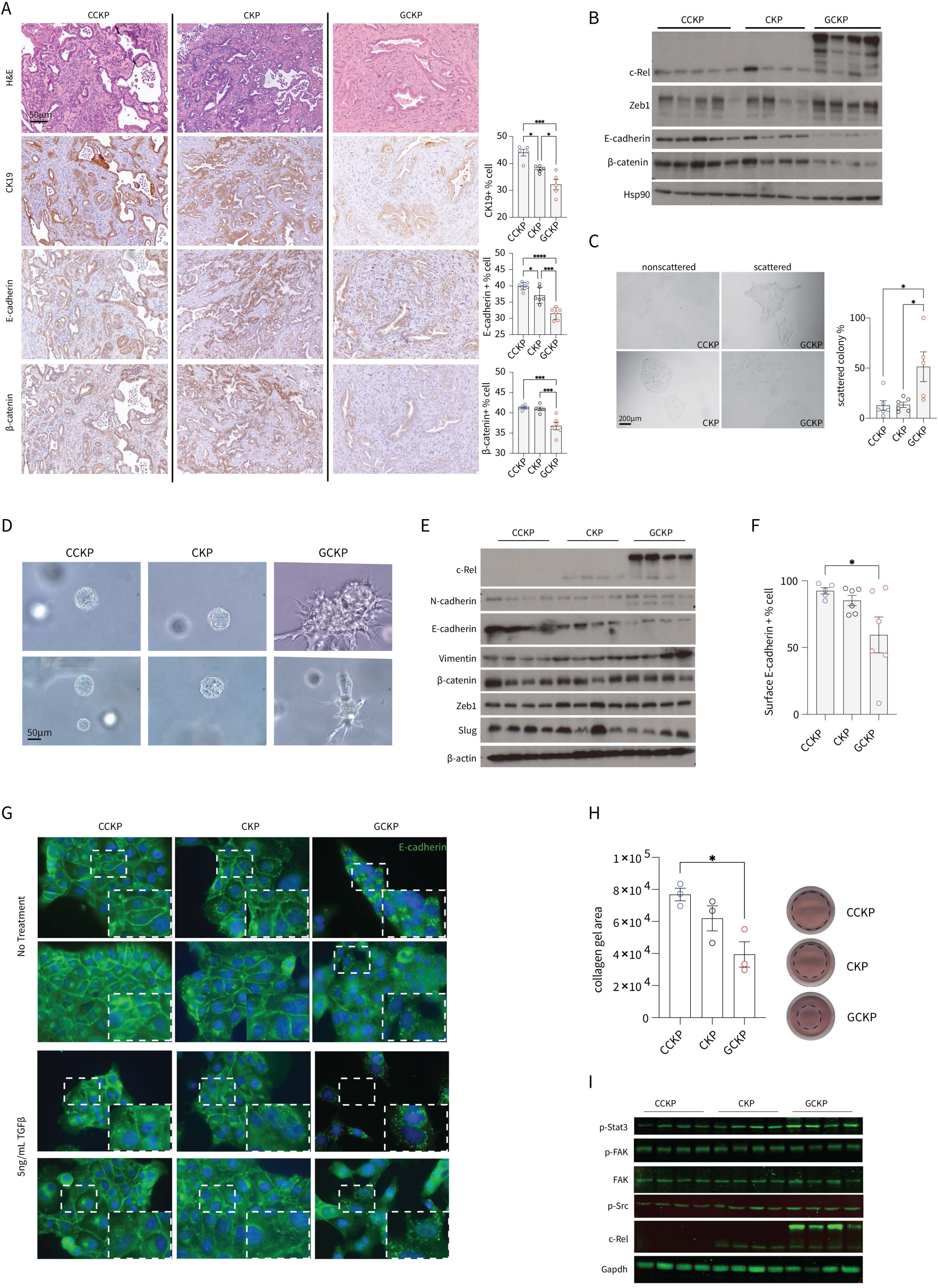
A) Representative H&E and IHC images for the respective targets and their quantifications. CK19 p= ***, E-cadherin p=****, and β-catenin p=*** B) Western blot analyses with bulk tumor protein lysates. HSP90 is used as a loading control. C) Representative scattered and non-scattered colony morphologies and their quantifications under a brightfield microscope (p=**). D) Representative images of organoids in matrigel from CCKP, CKP, and GCKP cells. E) Western blot analyses with lysates from isolated cancer cells at 70% confluency. β-Actin is used as a loading control. F) Surface E-cadherin expression as assessed by flow cytometry (p=*). G) E-cadherin immunofluorescence staining representative images are given. Cells are either not treated or daily treated with 5ng/mL TGFβ for two days (p=*) H) Collagen gels are visualized 4 days post-seeding (p=*). I) Western blot analyses with lysates from isolated cancer cells at 70% confluency under adherent conditions. Unless otherwise indicated, for all analyses ordinary one-way ANOVA test is used. ANOVA p values are given in the figure legends. Tukey’s multiple comparison tests are shown in the figures.

**Figure 3:**
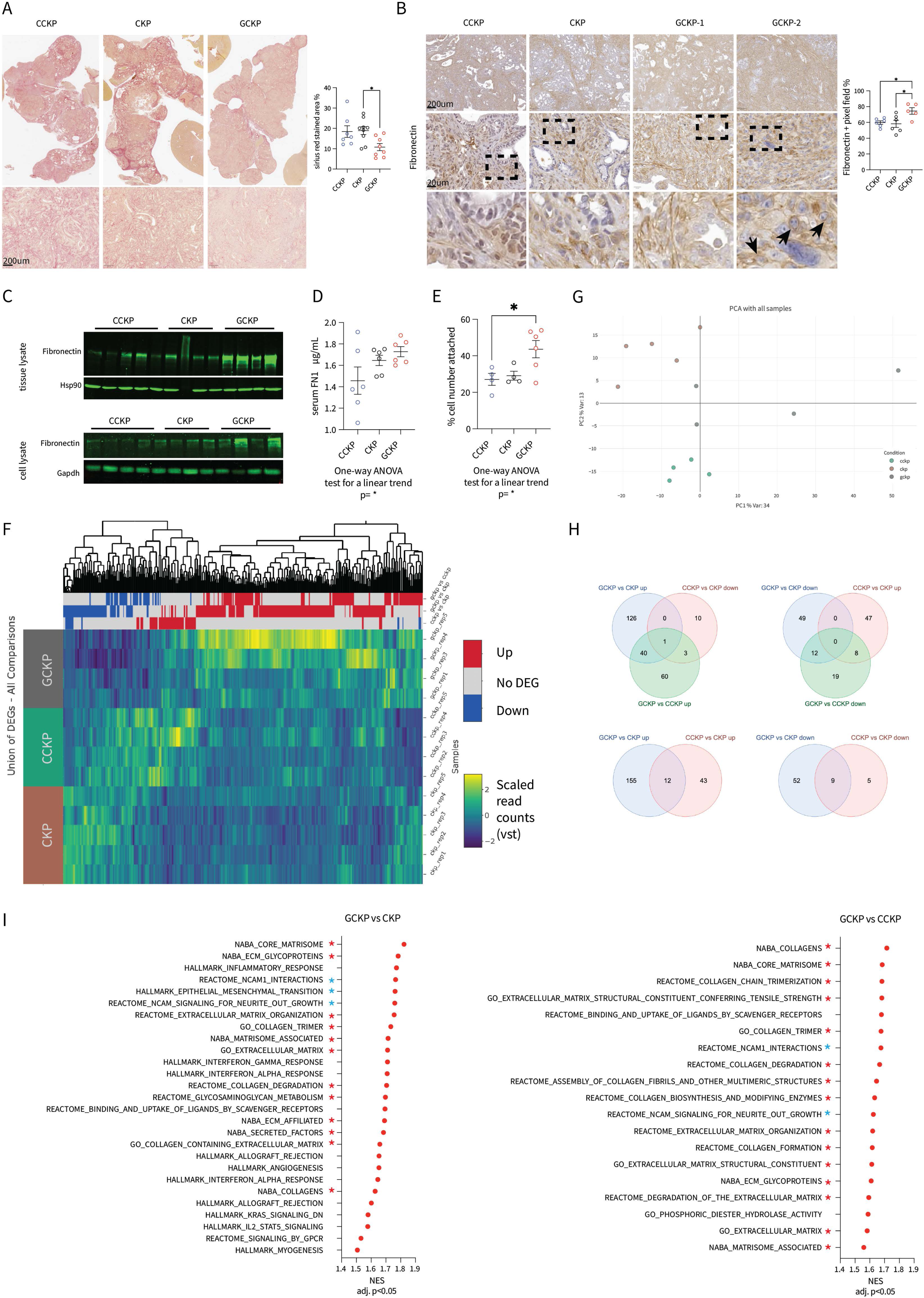
A) Representative images from Sirius-red stained tumors and their quantification (p=*) B) Fibronectin IHC representative images (p=*). GCKP-1 reflects FN1 deposition around cancer cells individually. GCKP-2 reflects perinuclear FN1 signal intracellularly. C) Western blot for Fibronectin and HSP90 as loading control on bulk tumor and isolated cancer cell lysates. D) Mouse serum FN1 ELISA results. The one-way ANOVA test for a linear trend is used (p=*). E) Fibronectin adhesion assay. The cell number attached to FN1-coated wells is normalized to the seeding number. The one-way ANOVA test for a linear trend is used (p=*). F) Heatmap for the differentially expressed genes of CCKP-CKP-GCKP isolated cells’ RNAseq analysis. G) Principal component analysis (PCA) plot of the cells used for the RNAseq analysis. H) Wenn diagrams illustrating the number of common genes up- or down-regulated in the given comparison pairs. I) GSEA analysis of the DEGs between given comparison pairs. Red stars mark enrichments related to ECM remodeling, while blue stars mark EMT-related pathways. Unless otherwise indicated, for all analyses ordinary one-way ANOVA test is used. ANOVA p values are given in the figure legends. Tukey’s multiple comparison tests are shown in the figures.

**Figure 4:**
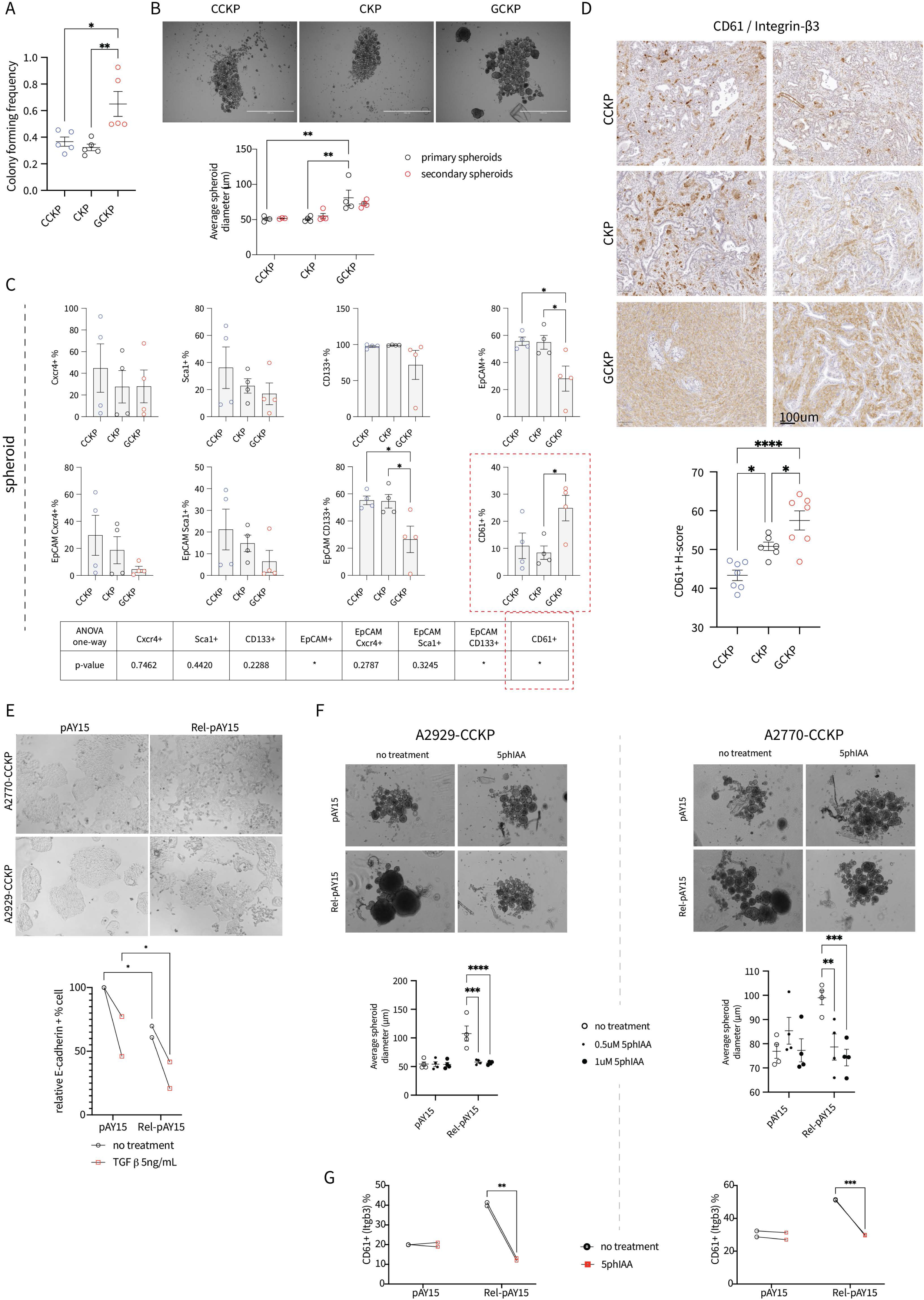
A) Colony-forming frequency is assessed by cells’ ability to form an organoid in matrigel (one-way ANOVA test p=**). Tukey’s multiple comparisons test results are shown in the figure B) Representative microscopic images of spheroids formed under non-adherent conditions. The average spheroid diameter is quantified and analyzed (genotype factor p=**). C) Surface expression of selected proteins on spheroids as assessed by flow cytometry analysis. p-values for the one-way ANOVA test are given in the table below. D) Representative IHC images for CD61 / Integrin-β3 and its quantification (one-way ANOVA test p=***). E) Representative microscopic images of CCKP cells stably transfected with either empty vector (pAY15) or c-Rel (Rel-pAY15) and treated with 5ng/mL TGFβ for two days. Relative surface expression of E-cadherin is analyzed by flow cytometry (c-Rel factor p=*). F) Representative microscopic images for degron cell spheroids. The left and right panels are for A2929 and A2770 CCKP cells respectively which are stably transfected. To rescue c-Rel expression from high to low, auxin analog 5phIAA is given in multiple doses. The average spheroid diameter is quantified and analyzed (genotype X auxin factors p= *** for both of the cell lines). G) Surface expression of CD61 is assessed by flow cytometry analysis of the spheroids (genotype X auxin factors p= ** and p=*** for A2929 and A2770 respectively. Unless otherwise indicated, for all analyses two-way ANOVA test is used. ANOVA p values are given in the figure or their legends. Šídák’s multiple comparison tests between genotypes are shown in the figures.

**Figure 5:**
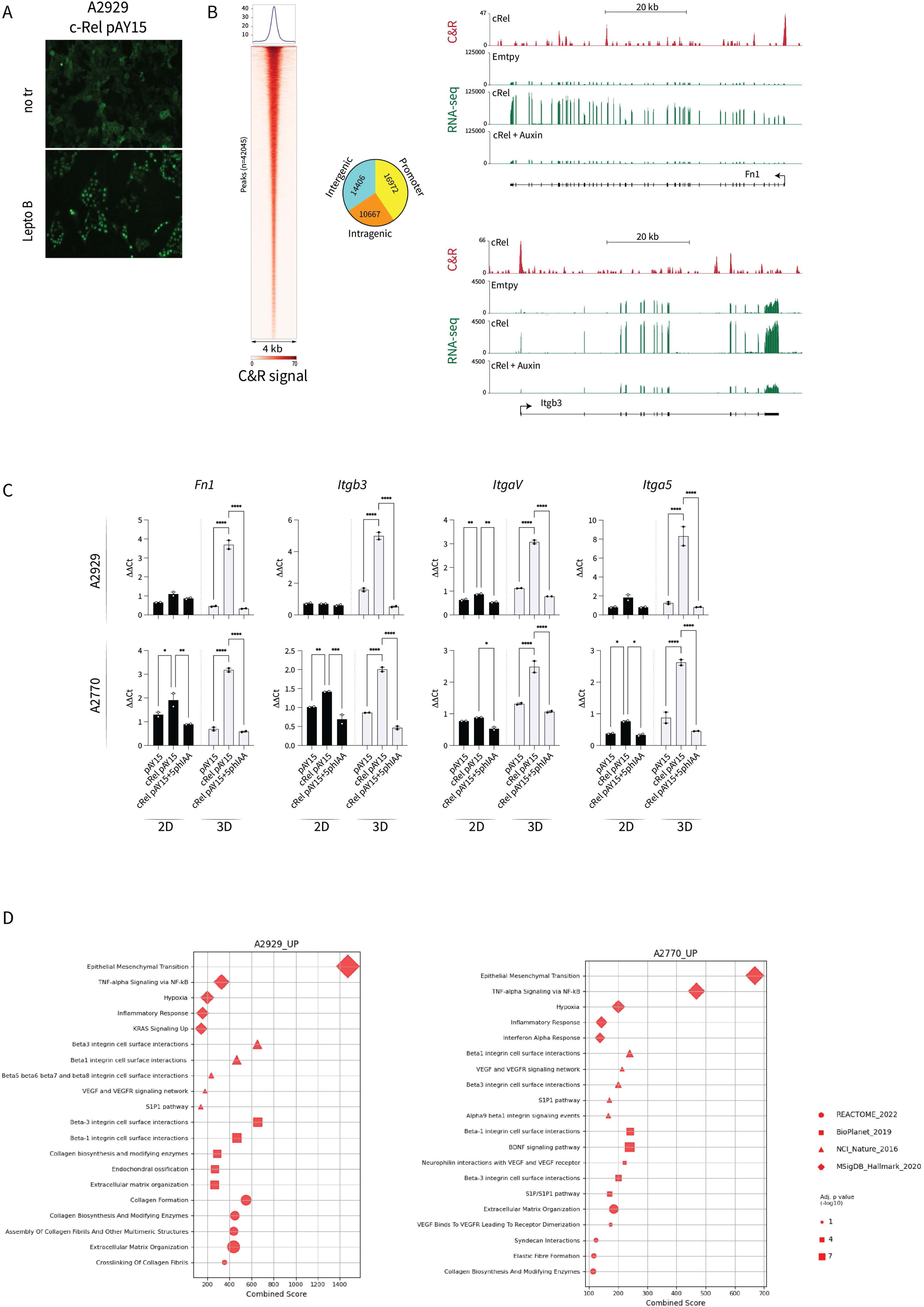
A) Representative fluorescent microscope imaging of the Rel-pAY15 stably transfected A2929 cell line. c-Rel is GFP tagged for tracing. Lepto-B: Leptomycin-B. B) Hetmap shows genomic regions differentially bound by c-Rel. Peak numbers for the genomic locations where c-Rel is bound are given as a pie chart. The CUT&RUN and RNAseq tracks display the expression levels of *Fn1* and *Itgb3* genes in mock-transfected, c-Rel overexpressed, and degron-induced c-Rel overexpressed A2929 cell lines. C) Normalized RT-qPCR analysis for the selected targets. Stably transfected cells are cultured either under adherent (2D) or non-adherent conditions to form spheroids (3D) (p values for A2929 c-Rel expression factor, *Fn1**, Itgb3***, ItgaV***, Itga5\*\**; for A2770 c-Rel expression factor *Fn1**, Itgb3**, ItgaV**, Itga5\*\**). D) Over-representation analysis performed by *Enrichr* for the Rel-pAY15 cells. The analysis is performed only with the genes which are differentially expressed by the Rel-pAY15 cells compared to pAY15 only, or Rel-pAY15 induced for degron. From each set, only the top 5 hits based on their combined scores are selected. Unless otherwise indicated, for all analyses two-way ANOVA test is used. ANOVA p values are given in the figure or their legends. Šídák’s multiple comparison tests between genotypes are shown in the figures.

**Figure 6:**
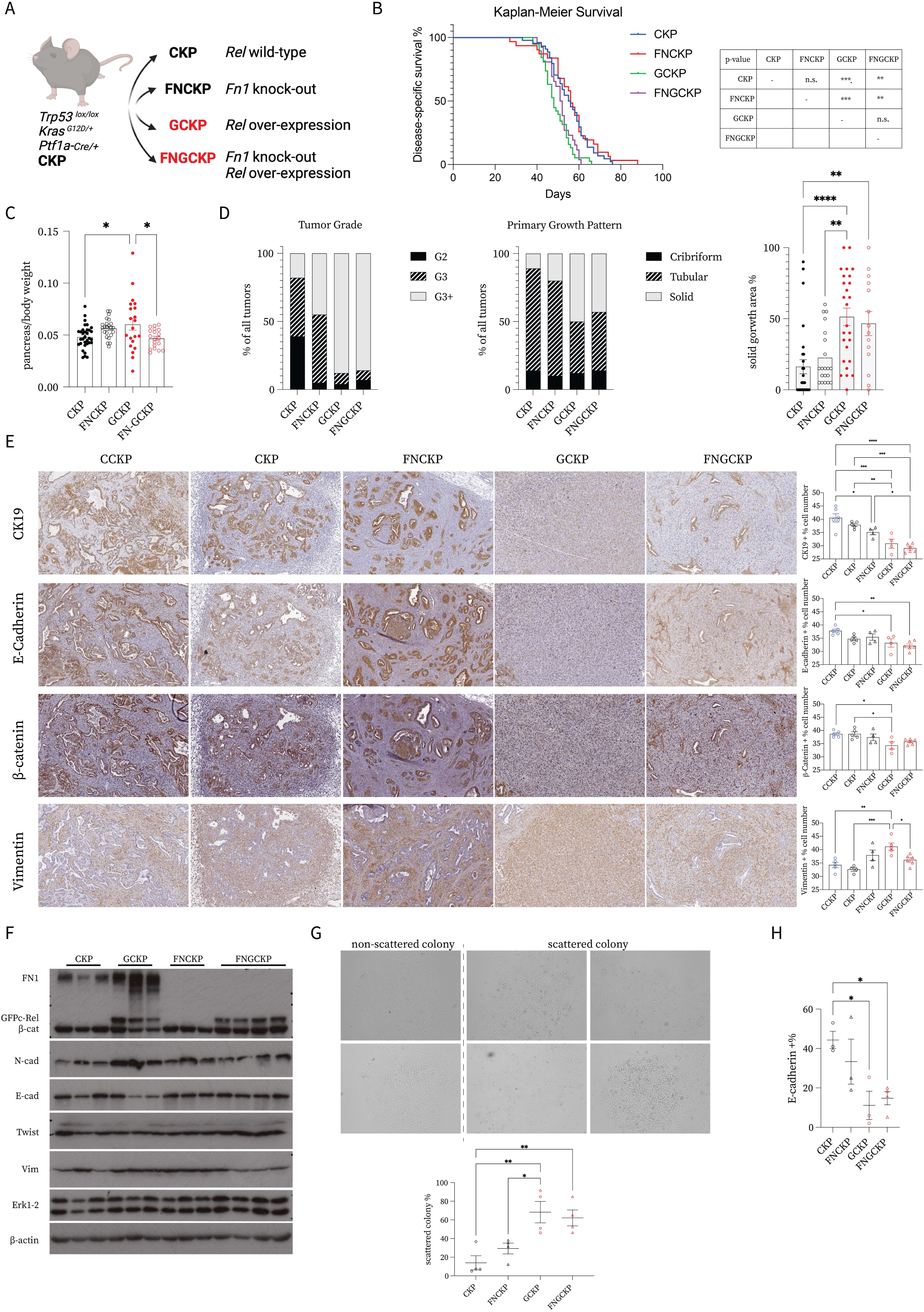
A) Schematic illustrating the compound mutant mice generation. B) Kaplan-Meier survival curves for the selected mice (CKP n=44, FNCKP n=31, GCKP n=38, FNGCKP n=26). Statistical analysis was performed with the Log-rank (Mantel-Cox) test in pairs and p values are displayed on the table. C) Pancreas/body weight comparison (p=**), D) Chi-square test (p=****) is used for tumor grade and primary growth pattern analysis. For the solid growth area % per tumor ordinary one-way ANOVA (p=****) and Tukey’s multiple comparisons test were used. The number of tumor mice analyzed is as given: CKP n=28, FNCKP n=20, GCKP n=26, FNGCKP n=14. E) Representative IHC images for the given EMT markers and their quantifications (CK19 p= ****, E-cadherin p=** and β-catenin p=**, Vimentin p=***). The scale bar is 100μm. F) Immunoblot analyses for some EMT markers with lysates from isolated cancer cells at 70% confluency. β-Actin is used as a loading control. Representative scattered and non-scattered colony morphologies and their quantifications under a brightfield microscope (p=**). H) Surface E-cadherin expression as assessed by flow cytometry (p=*). Unless otherwise indicated, for all analyses ordinary one-way ANOVA test is used. ANOVA p values are given in the figure legends. Tukey’s multiple comparison tests are shown in the figures.

**Figure 7:**
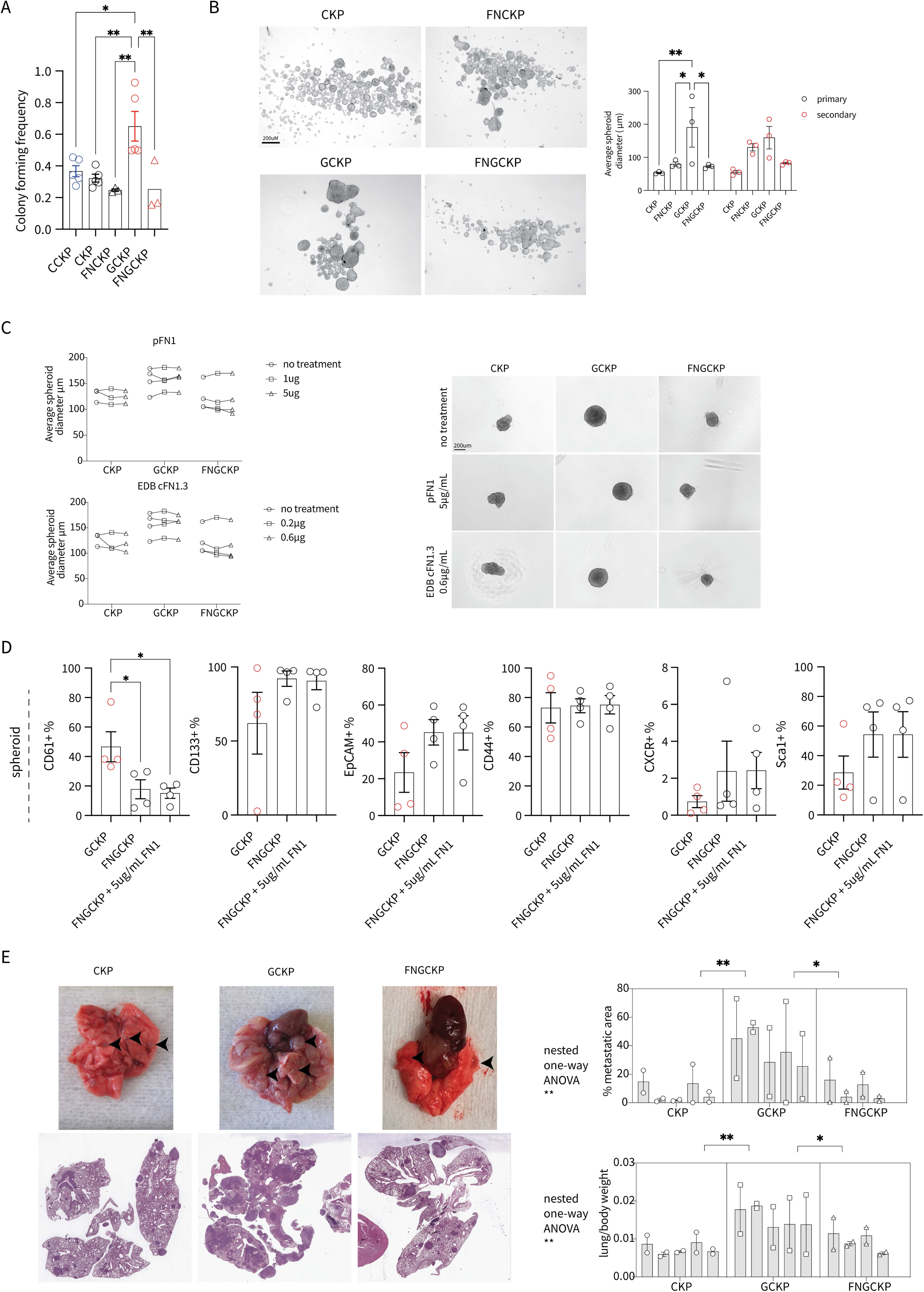
A) Colony-forming frequency is assessed by cells’ ability to form an organoid in matrigel (one-way ANOVA test p=**). Tukey’s multiple comparisons test results are shown in the figure. B) Representative microscopic images of spheroids formed under non-adherent conditions. The average spheroid diameter is quantified and analyzed (genotype factor p=*). C) Representative spheroid images and their diameter quantifications. The spheroid medium is supplemented with either human plasma fibronectin (pFN1) or cellular fibronectin with EDB domain + type II domains 8-14 (EDB cFN1.3) (treatment factor p= n.s.) D) Surface expression of selected proteins on spheroids as assessed by flow cytometry analysis (One-way ANOVA test CD61 p=*, CD133, EpCAM, CD44, CXCR4, Sca1 p= n.s.). Tukey’s multiple comparisons test results are shown in the figure. E) Representative macroscopic and microscopic images of the transplanted mouse lungs. Each cell line is injected two times. Nested one-way ANOVA is used for both of the analyses. Tukey’s multiple comparisons test results are shown in the figure. Unless otherwise indicated, for all analyses two-way ANOVA test is used. ANOVA p values are given in the figure or their legends. Šídák’s multiple comparison tests between genotypes are shown in the figures.

**Figure 8:**
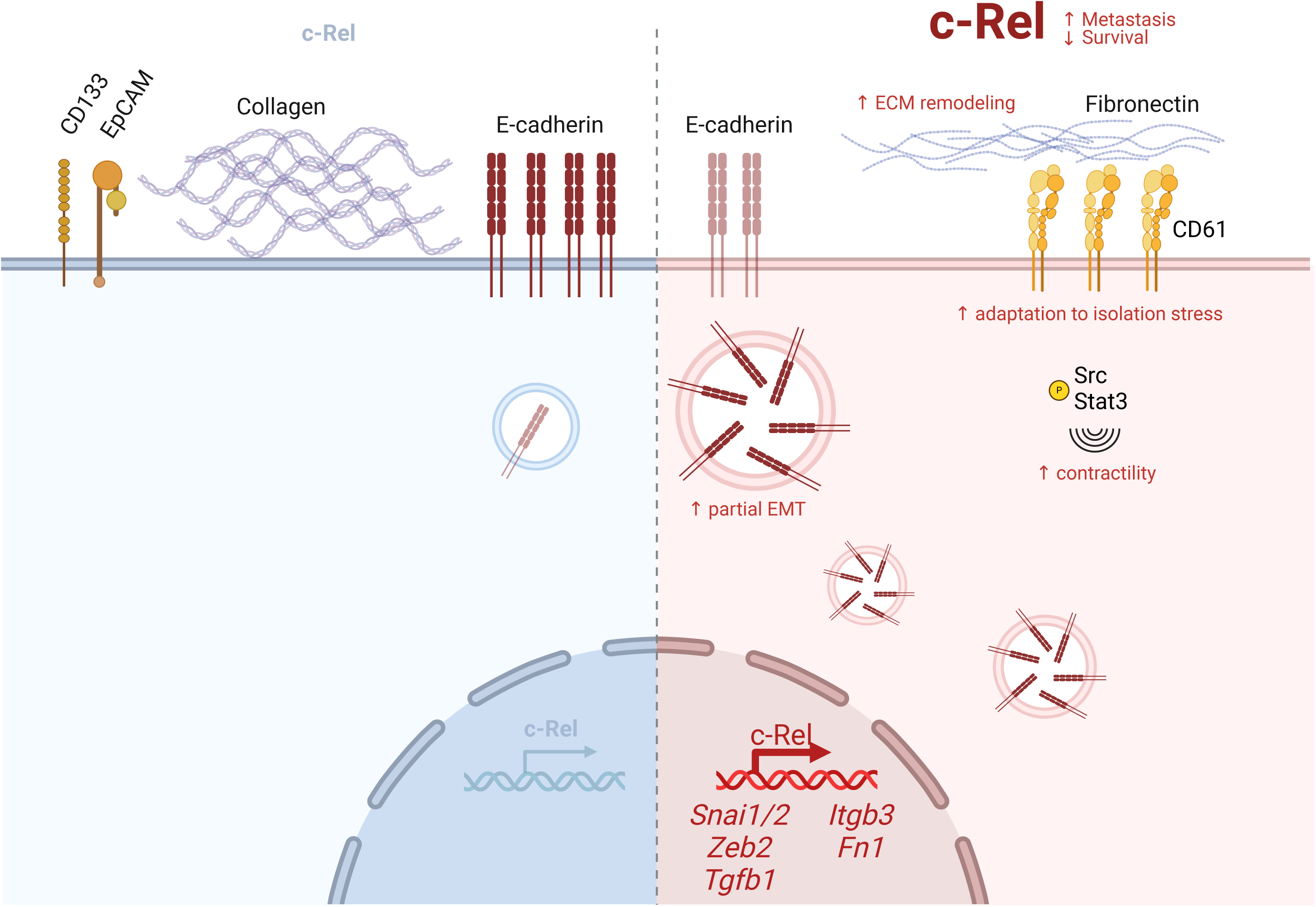
Schematic depicting the function of c-Rel in PDAC pathophysiology. c-Rel is an oncogenic factor in PDAC survival and metastasis. Increased c-Rel expression remodels ECM by converting collagen-rich stroma to fibronectin-rich. Cancer cells undergo epithelial to mesenchymal transition, increasing their plasticity (EMP) by reducing surface expression of E-cadherin and contractility via Src and Stat-3 phosphorylation. Once the isolation stress is induced under nonadherent conditions, c-Rel induces a niche enriched with fibronectin, coupled with higher CD61 expression. These changes are also supported with EMP, where c-Rel directly induces EMT-TFs like Snail, Slug, Zeb2, and TGFβ. Increased tolerance to isolation stress in the end drives metastasis. Created in BioRender. Kabacaoglu, D. (2025) https://BioRender.com/z35u460

## ACKNOWLEDGEMENTS

We thank to Elena Molina Roldan for her kind availability and cooperation with patients’ samples collection and along the TMAs’ processing. The development of the Bioinformatics for Next Generation Sequencing (BiNGS) shared resource facility is partially supported by the NCI P30 (P30CA196521) Cancer Center support grant, the ISMMS Skin Biology and Disease Resource-based Center NIAMS P30 support grant (AR079200), and the Black Family Stem Cell Institute. This work was also supported in part through the computational resources and staff expertise provided by Scientific Computing at the Icahn School of Medicine at Mount Sinai and supported by the Clinical and Translational Science Awards (CTSA) grant UL1TR004419 from the National Center for Advancing Translational Sciences. Research reported in this paper was supported by the Office of Research Infrastructure of the National Institutes of Health under award number S10OD026880. The content is solely the responsibility of the authors and does not necessarily represent the official views of the National Institutes of Health.

## METHODS

### 1) Mice

The genetically engineered mouse models used in the manuscript are previously described ^36^. Additionally, for the conditional deletion of fibronectin allele Fn1^tm1Ref^ model is used ^63^. In accordance with the European Directive 2010/63/EU, all animals are housed under specific pathogen-free conditions. The Federal German Guidelines for Ethical Treatment of Animals are followed during animal experimentation with approval (Regierung von Oberbayern). Mice are routinely checked for probable body recombination. The breeding is always maintained with fathers without *Ptf1a-Cre* to avoid germline recombination.

### 2) Experimental metastasis assay via intra-tail vein injection

Littermates without *Ptf1a-Cre* from the respective genotypes between 8-13 weeks old are used for the tail-vein injection of the mice. Cells are trypsinized, collected as single-cell suspension, and washed with PBS. Per mouse, 1 million cells are resuspended in 200μL PBS to be injected. Unless they show burden, mice are sacrificed 28 days after injection. To analyze metastatic colonization, the pancreas, liver, lung, spleen, duodenum, kidney, lung, heart, and thymus (if still present) are collected. Histopathological metastasis quantification is performed on the lungs. 2 -3 tissue sections per mouse are cut from the lungs with 150-200μm intervals to be stained for H&E, scanned, and quantified by QuPath ^64^. The metastatic colony and whole lung area are selected via the wand tool. Their ratio is assessed as a metastatic burden.

### 3) In vitro cell culture

#### a. Cell lines

Primary mouse pancreatic cancer cells are isolated from their respective mouse tumors. The cells are routinely cultured under standard 5% CO_2_, 37°C conditions, and tested for mycoplasma. The cell culture medium consisted of Dulbecco’s modified Eagle medium (DMEM)+ GlutaMAX (#10566016; Gibco) supplemented with 10%FBS (#10082147; Gibco) and 1% PenStrep (#1500-063; Gibco). Only cells under passage 10 are used for the subsequent experiments. Human pancreatic cancer cells are maintained based on ATCC recommendations.

#### b. Spheroid culture

Adherent cells are collected as single-cell suspensions after trypsinization. After cell quantification, all cells are serially diluted to 1000-2000 cells/mL per well to seed on Nunclon Sphera ultra-low attachment 24 well-plates (ThermoFisher Scientific 174930). The seeding number and choice of well-plates are altered based on the subsequent experiments. Cells are fed with spheroid medium when required. Primary spheroids are imaged 7-10 days post-seeding with a brightfield microscope. Spherical forms with diameters bigger than 40μm are quantified for size with Qupath software. Primary spheroids are trypsinized to form a single-cell suspension and reseeded into the spheroid medium again to obtain secondary spheroids. Spheroid culture is prepared as given: 0.5L DMEM/F12 (1:1) (Gibco, 21331-020), 5mL Pen/Strep (Gibco, 1500-063), 2mL Amphotericine B (R&D systems, B23192), 5mL L-Glutamine (Gibco, 25030-024), 10mL B-27 supplement (Gibco, 17504-044), 10μg FGFβ (R&D systems, 3718-FB).

#### c. Organoid formation assay in Matrigel

Tumor cells are trypsinized and serially diluted to obtain multiple cell densities as: 0.75, 3, 10, and 20 cells per 20μL in Matrigel per well. 7 technical replicates are seeded per density. After the Matrigel solidified, 80μL full medium is added to each well. 7 days post seeding, each well is evaluated under a microscope for spheroid formation (yes or no). The online software named ELDA: Extreme Limiting Dilution Analysis is used to quantify stem cell frequency ^65^.

### 4) RNAseq Analysis

The sub-confluent cancer cells are used for RNA extraction with the use of Maxwell® 16 LEV simplyRNA Purification Kit (Promega, AS1280) and Maxwell® 16 instrument (AS2000). RNA purity, concentration, and integrity are assessed by NanoDrop 2000 (Thermo Fisher Scientific) and gel electrophoresis. RNAs are then sent to Novogene Europe Company Limited (Cambridge, United Kingdom). Further RNA sample quality checks, mRNA library preparation with polyA enrichment, sequencing with NovaSeq 6000 PE150 platform, and quality control of RNAseq are performed by the company.

13 RNA-Seq libraries from three conditions (5 CKP, 4 CCKP, and 4 GCKP samples) systems are processed using the same pipeline for compatibility. Quality control has been performed using FastQC (v0.11.8) ^66^. Trim Galore! (version 0.6.5) has been used to trim the adapter sequences with a quality threshold of 20 ^67^. The mouse genome reference was GRCm38.p6 and GENCODE release M25 was used as the transcriptome reference. The alignment is performed by using STAR aligner (v2.7.5b) ^68^. Gene level read counts are obtained using Salmon (v1.2.1) for all libraries ^69^. All samples have passed the quality control requirements with > 90% of reads uniquely mapping (>20M uniquely mapped reads for each library) using STAR aligner.

Differential expression analysis was performed using the gene level read counts and the DESeq2 (v1.28.1) R package ^70^. Genes with less than 5 reads across all samples are filtered as inactive genes. A gene is considered differentially expressed if the adjusted p-value is less than 0.05 and the absolute log2 fold change is greater than 1. Gene set enrichment analysis for functional analysis is performed using the clusterProfiler R package (v3.16.0) ^71^. The gene sets used for functional analysis are obtained from The Molecular Signatures Database (MSigDB) ^72–74^. Gene expression heatmaps show the z-scores of DESeq2 VST normalized gene-level read counts and are generated using heatmaply R package (v1.1.0) ^75^.

For RNA-seq analysis of the degron cell spheroids, reads were aligned using STAR to the mm39 reference genome (settings: --quantMode GeneCounts --outSAMtype BAM SortedByCoordinate --outSAMunmapped Within --outSAMattributes Standard) ^68^. Subsequently, a count matrix file was generated from the STAR output. Differentially expressed genes were identified using the DESeq2 package with default settings ^70^, using cRel overexpressed cells as the reference sample. Genes with FDR < 0.01 and LFC > 1 or LFC < -1 were classified as differentially expressed. For visualization, BigWig files were generated with deeptools bamCoverage (settings: --binSize 1 --normalizeUsing RPKM) and tracks displayed on the UCSC Genome Browser. Heatmaps were created using the pheatmap package.

### 5) CUT&RUN analysis of the degron cells

A2929-cRel pAY15 stable transfectant cells are seeded to 6-well plates as 400.0000 cells per well. To induce c-Rel’s nuclear localization, cells are treated for 30 minutes with Leptomycin-B (20ng/mL, L2913, Sigma) in full medium. Before subsequent steps, c-Rel nuclear localization is assessed by visual GFP inspection. Required buffers are prepared and filtered through a 0.25 μm filter and stored at 4^°^C. <u>Buffer 1:</u> 5mL of 10X eBioscience Perm/Wash Buffer, 1 tablet of cOmplete EDTA free Protease inhibitor (Roche), 12.5 μL of 2M spermidine (molecular grade, Sigma), nuclease-free water up to 50mL. <u>Buffer 2:</u> 0.05% (w/v) molecular grade Saponin (25 mg in final 50mL, Sigma), 1 tablet of cOmplete EDTA free Protease inhibitor, 12.5 μL of 2M spermidine, up to 50mL PBS. <u>Antibody Buffer:</u> 40 μL of 0.05M EDTA, up to 10mL Buffer 1. <u>Calcium Buffer:</u> 20 μL of 1M CaCl_2_, up to 10mL Buffer 2. <u>2X Stop Buffer:</u> 200 μL of 0.5M EDTA, 40 μL of 0.5M EGTA, up to 5mL Buffer 2. The entire procedure is performed on ice, and all centrifugation steps are performed at 2000rpm for 4 min in cold. Cells are trypsinized and collected after 3X PBS wash to eliminate FBS-DNA on ice. Cells are incubated in 100 μL Antibody Buffer for 10 min and spun down. Cells are resuspended in 100 μL antibody buffer with c-Rel antibody (3 μg, AF-2699 R&D Systems), and incubated overnight on ice. Cells are washed 2X in 100 μL Buffer 1. Cells are resuspended in 50 μL Buffer 1 containing 1.5 μL pA/G-MNase (#40366, Cell Signaling), and incubated on ice for 1 h. Samples are washed 3X in 100 μL Buffer 2. For targeted digestion, cells are resuspended in 50 μL Calcium Buffer and incubated on ice for 30 min. 50 μL 2X Stop buffer is added to stop the reaction, and 1 μL of 0.1X Epicypher E. Coli spike in DNA (#40366, Cell Signaling) is added. The final mix is resuspended well and transferred to DNA low-binding eppies incubated at 37^°^C for 15 min and spun down for 5 min at 18.500g. 90 μL of the supernatant is carefully collected to subsequently purify DNA with the manufacturer’s protocol (MinElute, Qiagen). Finally, samples are eluted with 11 μL to be analysed for sample quality and sequencing by Novogene company.

All sequencing data underwent quality checks using FastQC. Cut&Run reads were aligned to the mm39 reference genome using Bowtie2 (settings: --local --very-sensitive)^76^. PCR duplicates and blacklisted regions were filtered out using Picard and Bedtools, respectively. Peaks were called using MACS2 (settings: -f BAMPE --call-summits --keep-dup all) ^77^. Deeptools was used to generate BigWig files (settings: --binSize 1 --normalizeUsing RPGC), and genomic tracks were visualized on the UCSC Genome Browser. Heatmaps were created with deeptools bamCoverage (settings: -a 2000 -b 2000 --skipZeros --missingDataAsZero --referencePoint center) ^78^. ChIPseeker was utilized for peak annotation ^79^.

### 6) Immunoblot analyses

Cells and tissues are collected in RIPA buffer (150mM NaCl, 1% NP-40, 0.5% Sodium deoxycholate, 50mM Tris pH:8.0) supplemented with fresh protease and phosphatase inhibitor cocktails (SERVA). Lysates are incubated on ice for 20 minutes and sonicated three times. Supernatant after centrifugation is used for Bradford Assay (Bio-Rad, 5000006) per manufacturer’s protocol. Lysates are diluted with 6X Laemmi buffer, and denatured at 95°C for 5 minutes, followed by immediate ice chill down. Lysates are then run on SDS-PAGE gel with varying concentrations depending on the target. The proteins are transferred to PVDF or NC membranes via the wet-transfer method. Membranes are incubated overnight with primary antibodies at 4°C on a shaker, and 1hr secondary antibody incubation. The target visualization is performed with ECL reagents or the IR-Dye LICOR system.

### 7) ELISA

Blood is collected from freshly sacrificed tumor mice. The serum is fractionated with Microvette® 500 Serum Gel CAT (Sarstedt, 20.1344) and snap frozen. The serums are diluted 1:8000 and used with a Mouse Fibronectin ELISA Kit (Novus, NBP2-60517). The values are log-transformed, and a standard curve is generated with linear regression. Based on the absorbance, the concentration for the unknown sample is calculated.

### 8) Tissue processing, histology quantifications, and immunohistochemistry (IHC)

Tissues are collected and fixed in 4% PFA for 48 hours. The tissues are then dehydrated and processed via Leica ASP300S. After paraffin embedding, 2μm tissue sections are rehydrated with 2 changes of Histoclear, 100%-96%-70% EtOH. The slides are evaluated based on H&E staining. For IHC, heat-induced antigen retrieval is performed either with citrate buffer (pH: 6), or Tris-EDTA buffer (pH: 8 or 9). Endogenous peroxidase is blocked with 3% H_2_O_2_. Blocking is performed 1hr with serum according to the secondary antibody species, and avidin (SP-2001 Vector). The primary antibody is diluted in a blocking solution with biotin and incubated overnight. DAB development is performed after secondary antibody and ABC-HRP incubation (Vector). Slides are then counterstained, dehydrated, mounted with Pertex, and scanned for subsequent digital pathological evaluation. For Sirius-Red staining, slides are deparaffinized, rehydrated, and stained with Weigert’s hematoxylin for 8 minutes, and Sirius-Red solution (0.5gr Sirius Red F3B (C.I. 35782) in 0.5L saturated picric acid solution) for 1 hour at RT. Slides are subsequently washed with 5% glacial acetic acid solution and water. Dehydrated and mounted slides are used for scanning. The IHC quantification is performed with Qupath software ^64^

### 9) Flow cytometry analyses

Cultured cells under adherent or spheroid culture conditions are trypsinized and suspended as single cells in FC buffer (PBS, 1% FBS, and 0.5mM EDTA). Then cells are incubated for 30min – 1hr with fluorophore conjugated antibodies for CSC cell surface expression markers. For compensation or unmixing, single stains of each antibody are used. Cells are finally resuspended in either 300μL PBS with 1:5000 Zombie-Red (BioLegend 423110), or 300μL FC buffer with 1:5000 DAPI (Cell Signaling, 4083S), depending on the panel. Before analyses, suspensions are passed through a 35μm strainer (Corning 352235). Flow run is performed with either Gallios Flow Cytometer (Beckman Coulter), Attune Flow Cytometer (Thermo Fisher Scientific), or Cytek Aurora Flow Cytometer. Downstream analyses are performed further with FlowJo software.

### 10) RT-qPCR

RNA is isolated as described in the RNAseq section. 600ng RNA is used for cDNA synthesis with a GoScript Reverse transcriptase kit (A2791, Promega) according to the manufacturer’s protocol. cDNA is diluted as 1:5 and 5μL is put for each PCR reaction using Go-Taq qPCR Master Mix (Promega). RT-qPCR is performed with either LightCycler-480 (Roche) or qTOWER^3^ G Cycler (Analytic Jena). The specificity of each primer pair is tested via gel electrophoresis and Tm calling.

### 11) Analysis of the publicly available datasets

#### a. Human PDAC transcriptomics analyses

The Bailey, Jandaghi, Janky, and TCGA datasets are used for GSEA analysis ^80–83^. The samples are ranked based on *REL* expression and divided into two groups from median as LOW vs HIGH. KEGG and HALLMARK geneset databases are used to perform GSEA via GSEA_4.3.2 ^72–74,84^.

#### b. pdacR stromal subtyping

The analyses are performed via the online tool pdacR (http://pdacr.bmi.stonybrook.edu/) ^34^. The datasets are filtered as given: TCGA-only whitelisted samples ^83^, Puleo-no prefiltering ^85^, CPTAC3-cellularity_call_from_VAF: Acceptable, sample_type: Primary Tumor ^86^. As gene sets to use the selected are as given: Moffit Normal 25, Activated 25, F13 _NormalStroma top100-top250, F5 _ActivatedStroma top100-top250 are used. As sample track, expression signature 1-user selected genes 1 is used, which is REL. Sample and Gene methods are selected as Pearson.

#### c. scRNA-seq data analysis

Previously published scRNA-Seq datasets GSE205013^87^, and GSE154778^88^, were retrieved from the NCBI GEO database. Barcodes, features, and matrix files were downloaded and imported into Python 3.9 using the Scanpy package (version 1.9.3) ^89^. Initial quality control was performed on each sample individually following best practices ^90^. Briefly, low-quality cells with mitochondrial gene content <15% were filtered out, along with genes present in fewer than 10 or 20 cells. Doublets were subsequently removed using the R package scDblFinder ^91^ with default settings. The datasets were then concatenated, and cells were further filtered based on the following criteria: fewer than 200 genes, fewer than 1500 counts per cell, or more than 150,000 counts per cell. Additionally, cells with >1% of transcripts representing erythroid genes were excluded. The data were normalized using the shifted logarithm method and log1p transformation. Dimensionality reduction was carried out on scaled data by computing PCA and a neighborhood graph. Batch correction and integration were performed using batch-balanced k-nearest neighbors (bbknn) (version 1.6.0) ^92^. Uniform Manifold Approximation and Projection (UMAP)^93^ was applied for two-dimensional visualization, and cells were clustered using the Leiden algorithm ^94^ with the default setting, Cell types were annotated using Scanpy’s rank_genes_groups function with default settings, method=’wilcoxon’, and corr_method=’bonferroni’. For downstream analysis, epithelial clusters were subset, and cells were stratified based on REL expression levels using a predefined threshold of 0.1, above and below which cells were classified as RELHigh vs RELLow respectively.

### 12) Auxin inducible degron 2 system (AID2)

pAY15 (OsTIR1(F74G) mAID-EGFP-NLS) was a gift from Masato Kanemaki (Addgene plasmid # 140534 ; http://n2t.net/addgene:140534 ; RRID:Addgene_140534) ^40^. To clone the mouse *Rel* gene, a PCR is conducted with cDNA generated from a GCKP cancer cell. The product and the vector are restriction digested by BamHI and MfeI, followed by ligation and transformation. The selected final clone is sequenced and validated. CCKP cells are transfected with either pAY15 (empty vector) or Rel-pAY15. Transient transfection was not suitable for degradation of c-Rel protein after 5phIAA treatment (varying doses and time points tried, 7392, Tocris). Based on personal communication with Dr. Kanemaki, pAY15, and Rel-pAY15 constructs are genome-integrated via transposase expression. Briefly, pCMV-TOL2 and pAY15 constructs are co-transfected to CCKP cells via polyplus JET Optimus, followed by puromycin selection (concentration is selected based on titration). pCMV-Tol2 was a gift from Stephen Ekker (Addgene plasmid # 31823 ; http://n2t.net/addgene:31823 ; RRID:Addgene_31823).

### 13) Immunocytochemistry

Cells are seeded to an 8-well chamber as 40.000 cells/well (94.6140.802, x-well, Sarstedt). After treatment, cells are washed with HBSS (Mg/Ca ++), and fixed at 37°C for 10 min with 4% PFA in HBSS. After 3x wash with HBSS, cells are permeabilized with 0.1% Triton-X in PBS for 5-7 minutes at RT. At 4°C, cells are incubated with primary antibody diluted in 2% filtered-BSA. The next day, after washing, cells are incubated with the respective secondary antibodies for 1hr at RT. After final washes, cells are mounted with DAPI (H-1200-10, Vector) and visualized with a Leica DMi8 microscope.

### 14) Collagen Contraction assay

Rat tail collagen Type I (9007-34-5, Corning) is diluted to 3mg/mL in sterile 0.1% acetic acid to prepare a working solution. Per cell line, 200.000 cells are prepared in 1mL medium, to be mixed with 0.5mL collagen working solution. For solidification of the collagen/cell culture mix, the NaOH amount is titrated. After NaOH addition, the solution is immediately aliquoted as 3×500 μL technical replicates to 24 well-plates. Each gel is carefully dissociated from the surrounding wall with a 10μL tip after the gel solidification. Gel contraction is observed for up to a few days until a difference is observed. Plates are scanned and gel surface areas are quantified by ImageJ software.

### 15) Fibronectin Adhesion assay

Human Fibronectin (1918-FN R&D Systems) is diluted as 1μg/mL in PBS supplemented with 0.5mM Mg^++^ and 0.9mM Ca^++^ and aliquoted as 100μL per well to 96wp Black/Clear bottom (165305, Thermo Fisher Scientific). The plates are incubated overnight at 4°C and next day blocked with 1% BSA-Fraction V in PBS for 30 min at RT. After the plates are slightly dried, 100.000 cells in 100μl PBS (supplemented with Mg^++^ Ca^++^) are seeded on wells with technical triplicates. Cells are spun down for 2 min at 300xrcf with a slow break. After 4 hours, unattached cells are discarded by 2X PBS wash. To normalize the seeding cell number, another group of cells is incubated for 24 hours without serum allowing all seeded cells to attach. Relative cell number is assessed via CyQuant viability assay (C7026, Thermo Fisher Scientific) per manufacturer’s protocols.

### 16) Statistics

Prism 9 software (GraphPad Software, Inc) is used for statistical analysis. Used p-values for all statistics are as given: ns p> 0.05, *p ≤ 0.05, **p ≤ 0.01, ***p ≤ 0.001, ****p≤ 0.0001). The data is displayed as mean ± standard error means (SEM).

### 17) Preparation of patient TMAs and their IHC

All patients who underwent duodenopancreatectomy from 2007 to 2013 at the Hepatobiliary and Pancreatic Surgery Unit (General and Digestive Tract Surgery Department, Clinico San Carlos Hospital) were initially assessed for eligibility, and samples were provided by its institutional Biobank (B.0000725; PT17/0015/0040; ISCIII-FEDER). The institutional review board (IRB) of University Hospital Clinico San Carlos evaluated the present study, granting approval on 10 March 2017 with approval number n° 17/091-E. The main criteria for eligibility were patients with a final histopathological diagnosis of localized pancreatic ductal adenocarcinoma (PDAC) who underwent surgery. Tumors were surgically resected and were formalin-fixed and paraffin-embedded immediately for pathologic diagnosis. To assess the survival analysis, only patients with a confirmed pathological diagnosis of PDAC and available data for progression-free and overall survival were included in the study.

#### a. Tissue Microarray

Tissue microarrays were constructed with 190 patients’ samples, 2 cores per patient, using the MTA-1 tissue arrayer (Beecher Instruments, Sun Prairie, WI, USA), with available formalin-fixed and paraffin-embedded tissues. A hollow needle was used to obtain a tissue core of 1 mm in diameter from selected tumor regions in formalin-fixed and paraffin-embedded tissues. These tissue cores were then inserted in a recipient paraffin block with precise spacing, resembling an array pattern. Sections from this paraffin block were then cut in a microtome and mounted on a microscope slide to be evaluated using immunohistochemistry.

#### b. Immunohistochemistry

Staining was conducted in 2 μm sections. Firstly, the slides were deparaffinized by incubation at 62 °C for 10 min and incubated for antigen retrieval using the PT-Link (Dako, Glostrup, Denmark) for 20 min at 97 °C in a low-pH buffered solution (EnVision Dako kit). The samples were then incubated with peroxide (EnVision Flex peroxidase blocking reagent) to block the endogenous peroxidase. The biopsies were stained overnight with diluted antibodies against c-Rel (1:50; R&D Systems AF-2699) followed by incubation with the appropriate anti-Ig conjugated with horseradish peroxidase (Anti-mouse/Anti-rabbit EnVision FLEX-HRP Labelled Polymer; Dako, Glostrup, Denmark) for 20 min. Visualization of the sections was carried out with 3,3′-diaminobenzidine (DAB, Dako, Glostrup, Denmark) as a chromogen for 5 min. Haematoxylin (Harris’ Haematoxylin, Sigma-Aldrich, San Luis, MO, USA) was used for counterstaining for 2 min.

All antibodies and anti-Ig horseradish peroxidase-conjugated antibodies presented high specificity, and no positiveness resulted from these antibodies individually. To determine the best immunohistochemistry conditions and maximize the specificity of antibodies, different healthy control tissues were used as positive controls according to “*The Human Protein Atlas*” (http://www.proteinatlas.org (accessed on 12 July 2022)). The evaluation of stainings has been performed by an expert pathologist (M.J.F.-A.).

#### c. Quantification of Immunohistochemistry

The evaluation of the immunoreactivity of the immunohistochemistry of different antibodies was carried out using a semiquantitative method called HistoScore, which considers the intensity of staining and the percentage of positively stained cells. The HistoScore (Hscore) ranged from 0–300 and multiplied the percentage of positively stained cells for each low, medium, or high staining intensity. The final score was determined using the following formula:

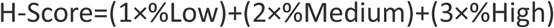

#### d. Ethical review board statement

The fundamental ethical principles promoted by the European Union have been strictly respected, as described in the Charter of Fundamental Rights of the EU (2000/C364/01). The project was evaluated by the Ethical Committee of University Hospital Clinico San Carlos (B.0000725; PT17/0015/0040; ISCIII-FEDER). The institutional review board (IRB) of University Hospital Clinico San Carlos evaluated the present study and approved it on March 10th, 2017, with approval number n° 17/091-E. All patients gave written informed consent for the use of their biological samples for research purposes. Moreover, the criteria of the Declaration of Helsinki (last revision 2013) and the Spanish Biomedical Research Law (14/2007, of July 3) were met. Ethical principles were followed regarding the use of human samples and generation of databases while respecting the Spanish Organic Law on Data Protection (LO 15/1999) and the European Union Fundamental Rights of the EU (2000/C364/01). The confidentiality of all personal, clinicopathological, and/or genetic data of patients will always be maintained per current legislation (2016/679 of the European Parliament and of the Council of April 27th, 2016 on Data Protection, Personal Data Protection Law 3/2018, of December 5th, on the Personal Data Protection that guarantee all digital rights, and the provisions in this regard contemplated in Law 41/2002, of November 14th.

## CONFLICTS OF INTEREST

The authors disclose no conflicts.

## FUNDING

This study was supported by the Deutsche Forschungsgemeinschaft (DFG, German Research Foundation) – mainly Project-ID 453134213, and partially Project-ID 456689823, Project-ID 492436553. DAR is supported by the German Research Foundation (DFG), CRC1479 (Project-ID:441891347-P17) and by the German Cancer Aid (grant number 70113697)

---

**Supplementary Figure 1.**
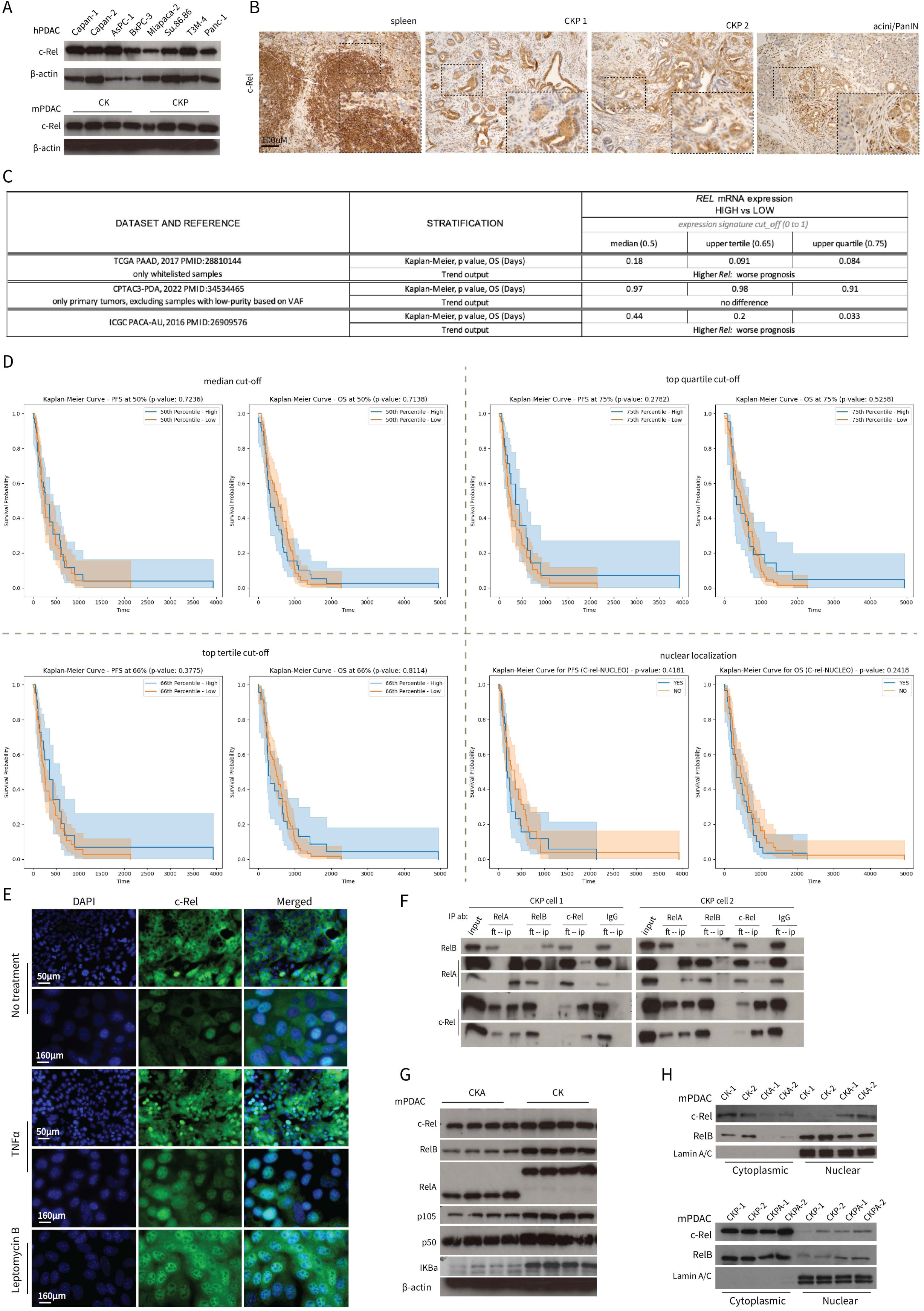

**Supplementary Figure 2.**
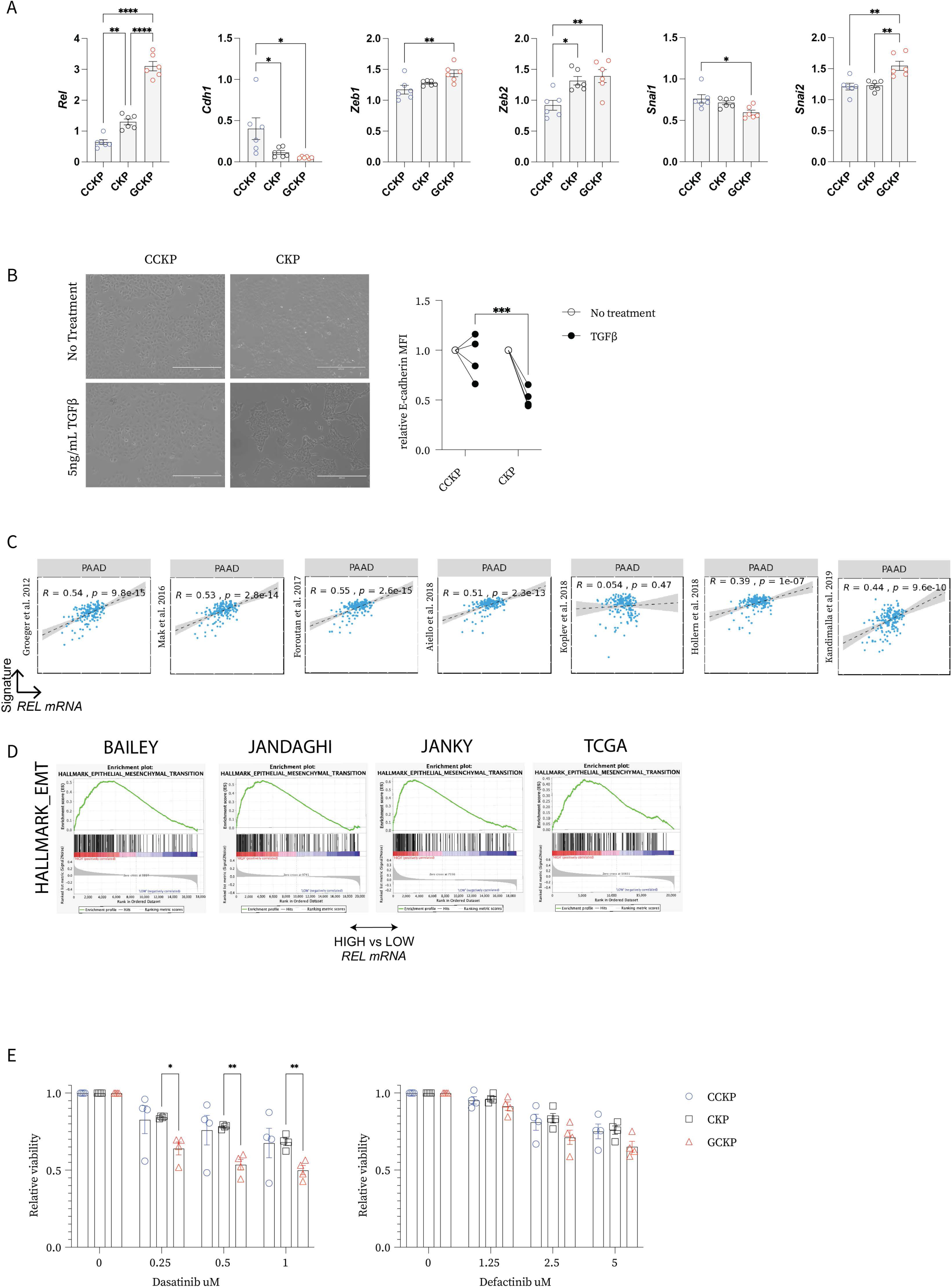

**Supplementary Figure 3.**
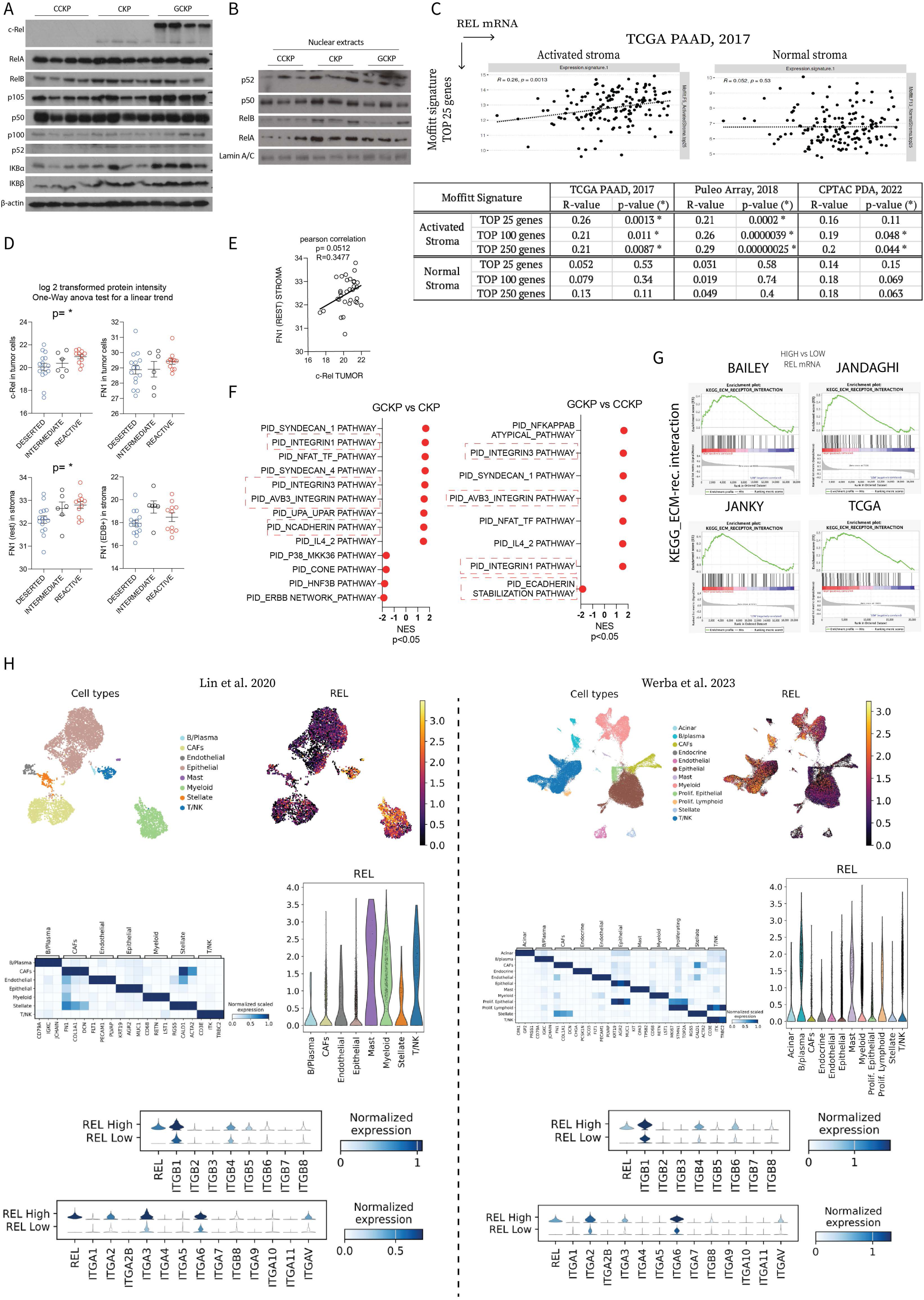

**Supplementary Figure 4.**
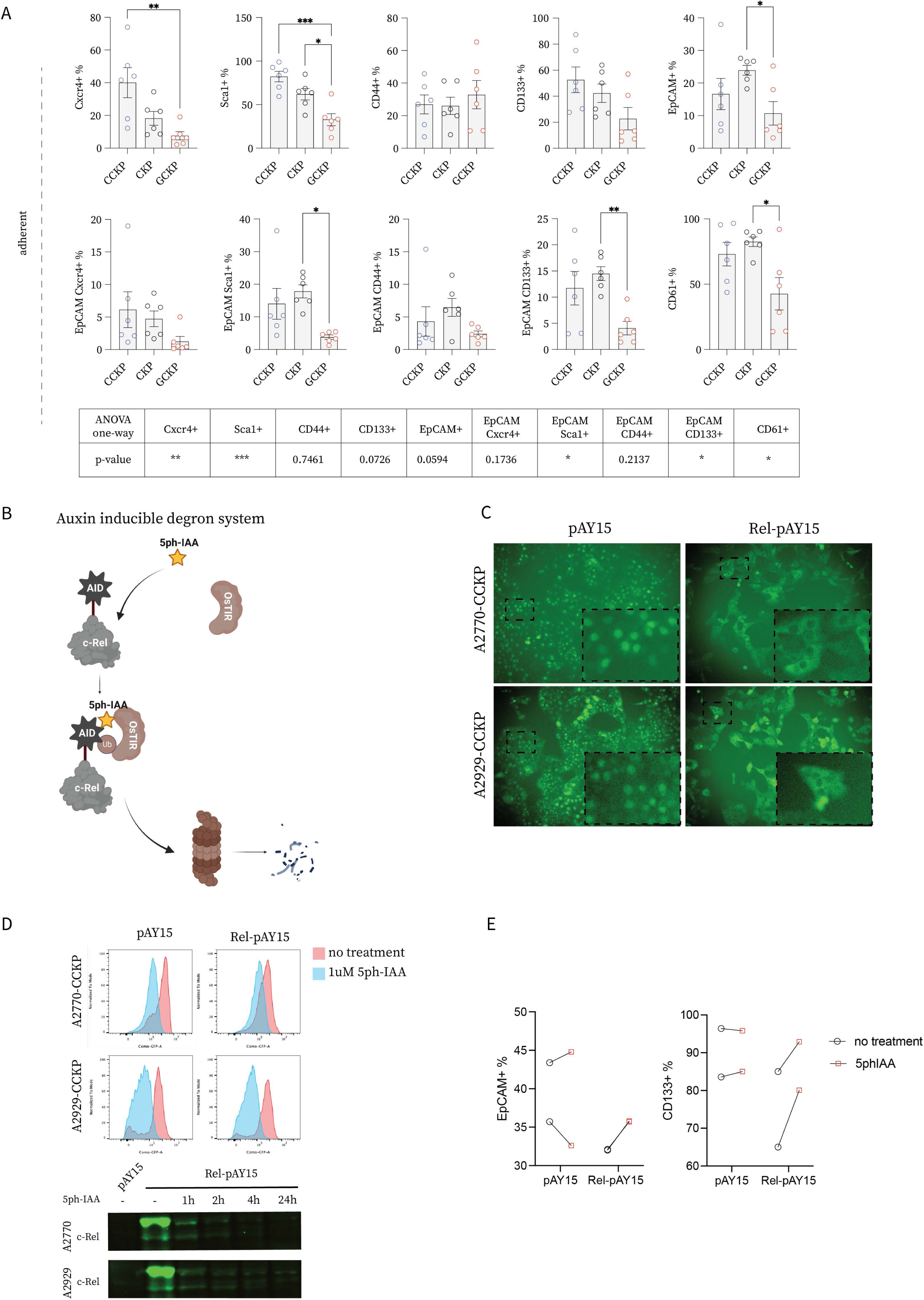

**Supplementary Figure 5.**
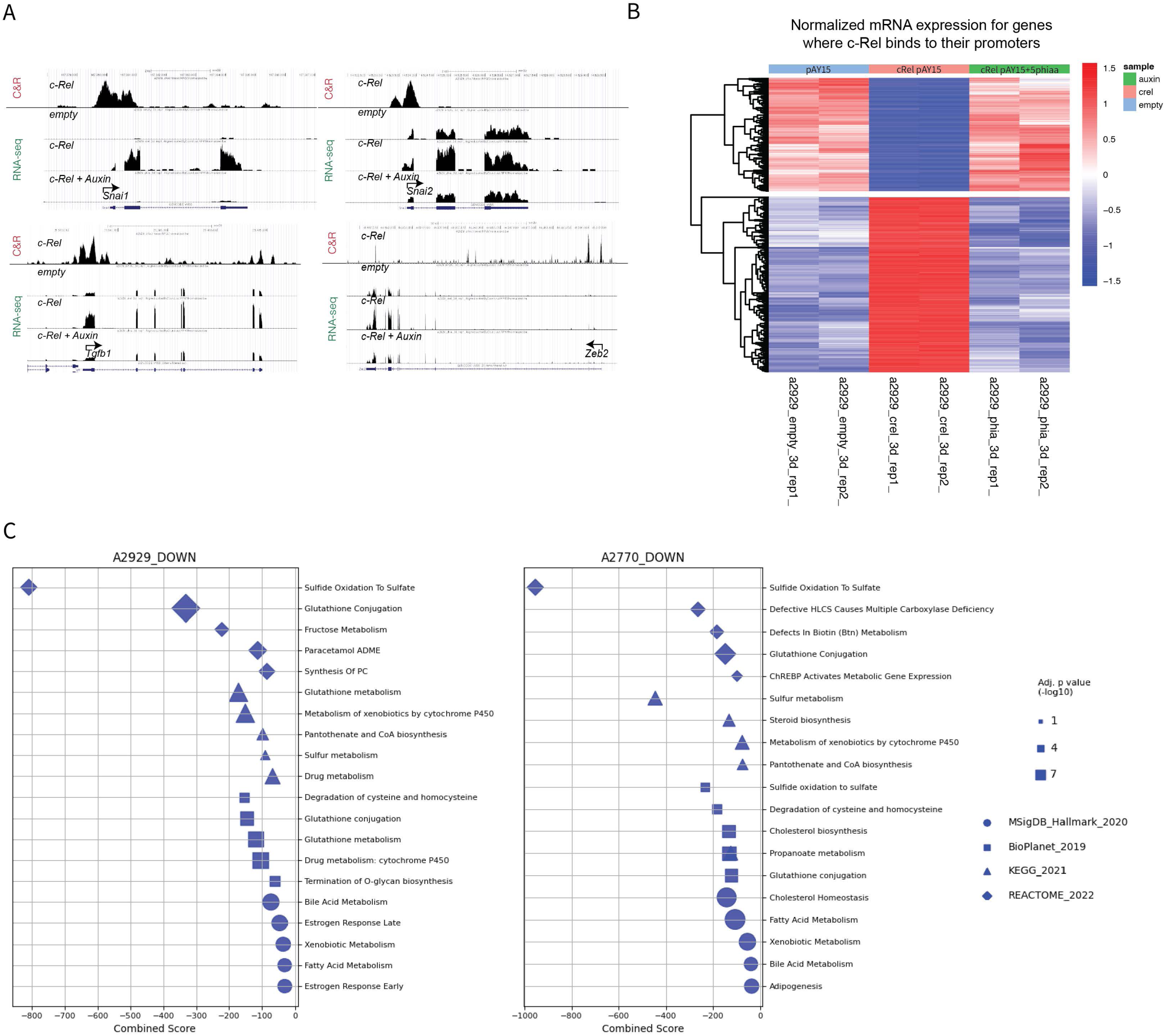

**Supplementary Figure 6.**
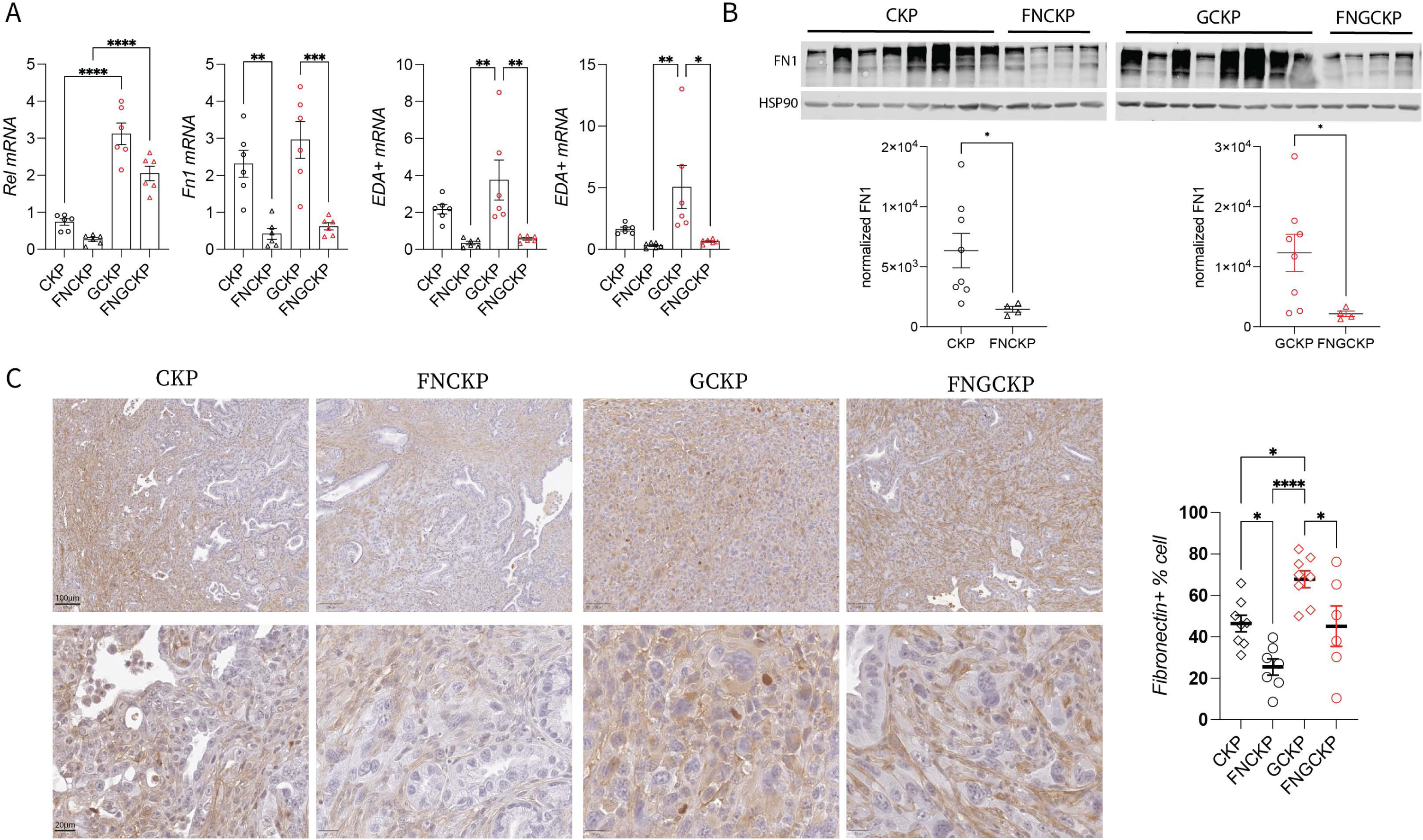

## Notes

### Competing Interest Statement

The authors have declared no competing interest.

